# A machine-learning tool to identify bistable states from calcium imaging data

**DOI:** 10.1101/2022.11.10.515941

**Authors:** Aalok Varma, Sathvik Udupa, Mohini Sengupta, Prasanta Kumar Ghosh, Vatsala Thirumalai

**Affiliations:** National Centre for Biological Sciences,Tata Institute of Fundamental Research, Bellary Road, Bangalore 560065; Department of Electrical Engineering, Indian Institute of Science, Bangalore - 560 012

**Author notes:** Current Address: Washington University School of Medicine in St Louis, 660 S Euclid Ave, St. Louis, MO 63110.

**Keywords:** Purkinje neuron, bistability, zebrafish, machine learning, convolutional neural network, recurrent neural networks, cerebellum

## Abstract

Mapping neuronal activation using calcium imaging *in vivo* during behavioral tasks has advanced our understanding of nervous system function. In almost all of these studies, calcium imaging is used to infer spike probabilities since action potentials activate voltage-gated calcium channels and increase intracellular calcium levels. However, neurons not only fire action potentials, but also convey information via intrinsic dynamics such as by generating bistable membrane potential states. While a number of tools for spike inference have been developed and are currently being used, no tool exists for converting calcium imaging signals to maps of cellular state in bistable neurons. Purkinje neurons (PNs) in the larval zebrafish cerebellum exhibit membrane potential bistability, firing either tonically or in bursts. Several studies have implicated the role of a population code in cerebellar function, with bistability adding an extra layer of complexity to this code. In this manuscript we develop a tool, CaMLSort which uses convolutional recurrent neural networks to classify calcium imaging traces as arising from either tonic or bursting cells. We validate this classifier using a number of different methods and find that it performs well on simulated event rasters as well as real biological data that it had not previously seen. Moreover, we find that CaMLsort generalizes to other bistable neurons, such as dopaminergic neurons in the ventral tegmental area of mice. Thus, this tool offers a new way of analyzing calcium imaging data from bistable neurons to understand how they participate in network computation and natural behaviors.

**Key Points Summary:** Calcium imaging – the gold standard of inferring neuronal activity – does not report cellular state in neurons that are bistable, such as Purkinje neurons in the cerebellum of larval zebrafish. We model the relationship between Purkinje neuron electrical activity and its corresponding calcium signal to compile a dataset of state-labelled simulated calcium signals.

We apply machine-learning methods to this dataset to develop a tool that can classify the state of a Purkinje neuron using only its calcium signal, which works well on real data even though it was trained only on simulated data.

CaMLsort also generalizes well to bistable neurons in a different brain region (ventral tegmental area) in a different model organism (mouse).

This tool offers a new way of analyzing calcium imaging data from populations of bistable neurons, thereby facilitating our understanding of how these neurons carry out their functions in a circuit.

## Introduction

Calcium imaging enables us to sample the activity of populations of neurons and even whole brains during animal behaviour (Ahrens et al., 2013; Grewe and Helmchen, 2009; Grienberger and Konnerth, 2012), providing us a window into the functioning of the nervous system. When a neuron fires an action potential, somatic calcium levels increase dramatically due to the opening of voltage-dependent calcium channels. Simultaneous electrophysiology and calcium imaging recordings of neuronal activity demonstrate the correspondence of somatic calcium imaging signals with action potential firing (Chen et al., 2013; Kerr et al., 2005; Kwan and Dan, 2012; Tian et al., 2009). Nevertheless, the calcium imaging signal is at best a delayed, low-pass filtered and non-linearly transformed proxy for neuronal activity. Non-linearities are introduced due to the dynamics of calcium release, and the kinetics of calcium binding and unbinding by the sensor molecules (Ali and Kwan, 2020). Yet, this technique is useful in cases where the neurons being studied have low baseline firing rate. However, this is not often the case and several neuronal classes, such as cerebellar Purkinje neurons and cortical interneurons are known to fire at very high spike rates spontaneously (Armstrong and Edgley, 1984; Markram et al., 2004). Further, many neuronal types have interesting intracellular dynamics apart from action potential firing that might strongly modulate calcium imaging signals (Lin et al., 2007; Moreaux and Laurent, 2007). Lastly, confounds may be introduced when more than one type of electrical event leads to somatic calcium increase (Lev-Ram et al., 1992; Ramirez and Stell, 2016).

Purkinje neurons (PNs) show all of the features mentioned above that make the interpretation of calcium signals in these neurons especially difficult. PNs are principal neurons of the cerebellum, integrate thousands of synaptic inputs and are indispensable for cerebellar function (Narayanan and Thirumalai, 2019). Therefore, studying PN population activity during motor behaviours is critical to deciphering cerebellar computation. PNs fire simple spikes and complex spikes - in mammals, climbing fiber (CF) input from the inferior olive leads to calcium entry within the dendrites and several spikelets that ride the depolarization wave (Kitamura and Häusser, 2011; Miyakawa et al., 1992); in fish, the olivary input causes an all-or-none AMPAR-mediated giant synaptic potential with concomitant calcium entry but without any spikelets (Sengupta and Thirumalai, 2015). While the CF input causes a relatively large calcium signal, bursts of simple spikes can also generate calcium signals via the activation of somatic voltage-dependent calcium channels, albeit smaller in amplitude (Knogler et al., 2019; Sengupta, 2015). This means that optically recorded calcium dF/F signals cannot be uniquely assigned to simple spikes or CF inputs.

In addition to the above, larval zebrafish PNs exhibit membrane potential bistability – these neurons can exist in one of two stable membrane potentials. At a depolarized state, they generate action potentials at a more or less regular rate and are said to be tonic. When hyperpolarized, they generate bursts of action potentials (Sengupta and Thirumalai, 2015). Simple spikes in the bursting state are highly correlated with motor bouts whereas those in the tonic state are not (Sengupta, 2015). This implies that bistability has important ramifications for how motor bouts are represented across the entire PN population. We wondered if calcium imaging can be used to deduce the state of PNs. Since both CF inputs and simple spikes contribute to calcium signals, assigning optically recorded dF/F signals is not straight-forward and deduction of spike rate using existing spike inference algorithms is not possible.

Here, we first performed simultaneous electrophysiological recordings and calcium imaging in larval zebrafish PNs to obtain ground truth data. Unsurprisingly, we found that none of the usually measured parameters such as peak amplitude, area under the curve etc., of the calcium signal uniquely correlated with cellular state. We therefore built a sorter using convolutional recurrent neural networks to identify cellular state from dF/F data. We have tested our sorter with data that was not part of the training set and validated that it performs at above 90% accuracy and with F1 scores greater than 0.8 in most cases. Our method demonstrates that calcium imaging datasets can be mined using such tools to glean greater insights into neuronal activity states.

## Materials and Methods

### Ethical approval

All experimental procedures were approved by the institutional animal ethics committee (IAEC) and institutional biosafety committee (IBSC) at the National Centre for Biological Sciences.

### Experimental Animals

Experiments were performed on Indian wild type (Ind WT) zebrafish (*Danio rerio*). Embryos were obtained by setting up an in-cross between adult Ind WT zebrafish raised and housed in a ZebTec multi linking system (Tecniplast, Italy) with a pH setting of 7.8 and conductivity of 1200μS. Larvae were kept in a MIR-154 incubator (Sanyo, Japan) with 14:10h light-dark cycle maintained at 28°C in E3 medium (composition in mM: 5 NaCl, 0.17 KCl, 0.33 CaCl_2_,and 0.33 MgSO4, pH 7.8) in standard 90mm Petri dishes (Tarsons, Kolkata, India). The medium was routinely replaced, and starting at 5 dpf, larvae were fed Zeigler Larval diet AP100 (<100 microns) (Pentair AES, FL, United States). Larvae have not yet undergone sex specification by these stages, so experiments performed were agnostic to the sex of the animal.

### Transient transgenesis by microinjection

Thin-walled borosilicate capillaries (1.0mm OD, 0.56mm ID; Warner Instruments, Hamden, CT, United States) were pulled using a Flaming-Brown P-97 pipette puller (Sutter Instruments, Novato, CA, United States) to produce long and thin pipettes for microinjection. Indian WT embryos at the 1-2 cell stage were injected with ∼10nL of a mixture of tagRFP-T:PC:GCaMP5G plasmid (Matsui et al., 2014) (50 ng/μL) and Tol2 transposase mRNA (50 ng/μl) (Urasaki et al., 2006), using a Picospritzer III (Parker Hannifin, NH, United States) pressure injection apparatus, and raised as described above. Larvae were screened at 5 dpf at an epifluorescence dissection microscope (Olympus SZX-16) and those showing mosaic labelling of Purkinje neurons were selected for experiments, which were performed at 7 dpf at room temperature (25-28°C).

### Electrophysiology

Larvae were anaesthetised in 0.01% MS-222 (Sigma-Aldrich; Missouri, USA) and transferred to a custom-made acrylic recording chamber. Fine tungsten wire (California Fine Wire, CA, USA) was used to pin the larva onto a piece of Sylgard (Dow Corning, Midland, MI, United States) glued to the recording chamber. Two pins were put through the notochord at the level of the swim bladder and the tail, respectively. A third pin was then used to position the larval head dorsal-up. The MS222 was replaced by external solution (composition in mM: 134 NaCl, 2.9 KCl, 1.2 MgCl_2_, 10 HEPES, 10 glucose, 2.1 CaCl_2_, 0.01 D-tubocurarine; pH 7.8; 290 mOsm) and skin on the dorsal surface of the larva was carefully removed using fine no. 5 forceps (Fine Science Tools, Foster City, USA) to expose the brain.

A Flaming-Brown P-97 pipette puller (Sutter Instruments, Novato, CA, United States) was used to pull patch pipettes made of standard-walled borosilicate capillaries with a filament (1.5 mm OD; 0.86 mm ID; Warner Instruments, Hamden, CT, United States), pulled to a final tip diameter of 1-1.5 μm. For loose patch recordings, pipettes were backfilled with external solution, while for whole-cell recordings, pipettes were backfilled with K-gluconate internal solution (composition in mM: 115 K gluconate, 15 KCl, 2 MgCl_2_, 10 HEPES, 10 EGTA, 4 Mg-ATP; pH 7.2; 290 mOsm).

Solutions were filtered using a 0.2μm filter (Millipore, Merck, Germany) before backfilling pipettes, which typically had resistances between 8 and 10 MΩ. Sulforhodamine (Sigma-Aldrich, St.Louis, MO, United States) was added to the patch internal solution at a final concentration of 5μg/mL to facilitate visualization of cellular morphology post-recording.

Recordings were acquired with a Multiclamp 700B amplifier, Digidata 1440A digitizer and pCLAMP 10 software (Molecular Devices, Sunnyvale, CA, United States). Data were low pass filtered at 2 kHz and sampled at 20-50 kHz with a gain of 1 (whole-cell recordings) or between 20-1000 (loose patch recordings), so as to optimally use the dynamic range of the digitizer.

### Simultaneous calcium imaging and electrophysiology

Indian WT larvae microinjected with the Tol2-tagRFP-T:PC:GCaMP5G construct (gift from Dr. Hideaki Matsui, Niigata University, Japan) and screened for sparse labelling were used for this experiment. Purkinje neurons expressing GCaMP5G were targeted for the recordings. Widefield imaging was performed using a 60x water-immersion objective (1.0 NA) at an Olympus BX61WI microscope. Excitation light was provided using a pE300-ultra (CoolLED, Andover, England). Imaging was performed using an EVOLV EM-CCD camera (Photometrics, Tucson, AZ, United States) at the rig, with an exposure of 30 ms, and an on-chip gain of 170. An ROI was drawn around the targeted cell, so that only an isolated cell was imaged.

Imaging was synchronized with electrophysiology by interfacing with the Digidata 1440A.

### Calcium Imaging Analysis

Image analysis was performed in Fiji/ImageJ (Schindelin et al., 2012). In order to identify cell boundaries, the averaged time series was used as a reference and an ROI drawn around the cell. Bleach correction was performed using the Exponential Fitting Method in the Bleach Correction plugin in Fiji. The average raw pixel intensity values within the ROI for each time point was extracted. The baseline intensity, F0, was taken to be the 5th percentile of these values. dF/F was then calculated for each frame using the formula (F - F0)/F0, where F represents the fluorescence in that frame and F0 the baseline fluorescence. Following this calculation, a Gaussian filter was applied to the time series in MATLAB R2019b (Mathworks, Natick, MA), using its function ‘*smoothdata’*, with a window size of 15.

To detect transients in the calcium signal, dF/F traces were first normalised to a range of [0, 1] using min-max feature scaling. Peaks in the normalised trace were then detected using the ‘*findpeaks’* function in MATLAB, with a minimum peak height of 0.1, a minimum peak prominence of 0.05 and a minimum interval of 500ms between consecutive peaks.

### Spike detection and analysis

All electrophysiological analyses were done using custom scripts written in MATLAB, using functions defined in a publicly available repository (https://github.com/wagenadl/mbl-nsb-toolbox) to read electrophysiology data. For extracellular recordings, spikes were identified as events that showed large fluctuations relative to a 25ms rolling window baseline and the peak timing and amplitude were extracted. Events were sorted based on amplitude, with small amplitude events being classified as simple spikes and large amplitude events as climbing fiber (CF) inputs, as per (Sengupta and Thirumalai, 2015). For intracellular recordings, peak detection algorithms inbuilt in MATLAB were used to identify spikes. Once again, events were sorted as simple spikes or CF inputs based on their amplitude relative to baseline.

### Calcium imaging reconstruction from electrophysiology data

Simultaneous calcium imaging and extracellular electrophysiology data was used to infer optimal GCaMP5G kernels, which are of the general form

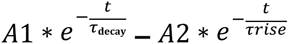

for simple spikes and CF inputs of zebrafish PNs. Simple spike and CF input timings were first extracted from the electrophysiology data for a single cell. Simple spike timings were convolved with a simple spike kernel, with amplitudes A1s and A2s and CF input timings were convolved with a similar CF input kernel, with amplitudes A1c and A2c. The sum of both these convolutions was taken to be the putative calcium imaging reconstruction, down sampled to match the sampling rate of calcium imaging. The mean squared error (MSE) between the reconstruction and the true calcium imaging trace was calculated. The values for A1s, A2s, A1c and A2c were obtained by minimizing the MSE between the true calcium imaging trace and the reconstructed trace using gradient descent. To ensure that local minima were avoided, the gradient descent algorithm was repeated with randomized starting points. The kernels with these coefficients were taken to be the optimal simple spike and CF input kernels, respectively, for that cell.

The ‘optimal’ kernel would be one that generalized best. This was tested by using the kernels from one cell to reconstruct calcium imaging for other cells for which ground truth data was available. As before, the MSE and also the Pearson correlation coefficient between the reconstructed trace and the true trace were calculated. The final ‘optimal’ kernel was obtained by taking the kernel which generalized the best, i.e., had minimum MSE and maximum correlation across all cells. To test whether the correlation was real or not, the Pearson correlation coefficient between a scrambled reconstructed trace and the true trace was also calculated, for reference.

This process of convolving simple spike and CF timings with their optimal GCaMP5G kernels and summating the resultant traces was then used to generate a calcium imaging trace from both extracellular and intracellular Purkinje cell recordings.

### Building a state-labelled calcium imaging dataset

Intracellular and extracellular recordings from larval zebrafish Purkinje neurons acquired during the period 2011-2020 by various members of the lab were compiled. Good quality recordings were those in which the resting membrane potential of the cell was not below -80mV or not above -20mV, were uninterrupted gap-free recordings with no current injections and had two clearly distinguishable event types. Besides poor quality recordings, those recordings with an ambiguous, indeterminate state were also excluded. Only those recordings which met the aforementioned criteria were retained. Then, recordings were renumbered randomly, to remove grouping of data collected by the same individual. Recordings were then classified as tonic or bursting, first by visual inspection. These classifications were then further verified by inspecting the inter-event intervals of simple spikes. While inter-event intervals from tonic cells form a single cluster, bursting cells have inter-event intervals that trail to large values, corresponding to the inter-burst intervals. Lastly, the coefficient of variation of the inter-spike intervals was used as an extra parameter of interest. These three pieces of information were used in conjunction (**Fig. 1D-G**), but the final classification relied on visual inspection. If there was a switch in state within the recording, the recordingswere labelled as “switch”, with the appropriate state switch timing and kind (tonic-to-bursting, or bursting-to-tonic) marked. A summary of the recording information is in Table 1.

**Figure 1.**
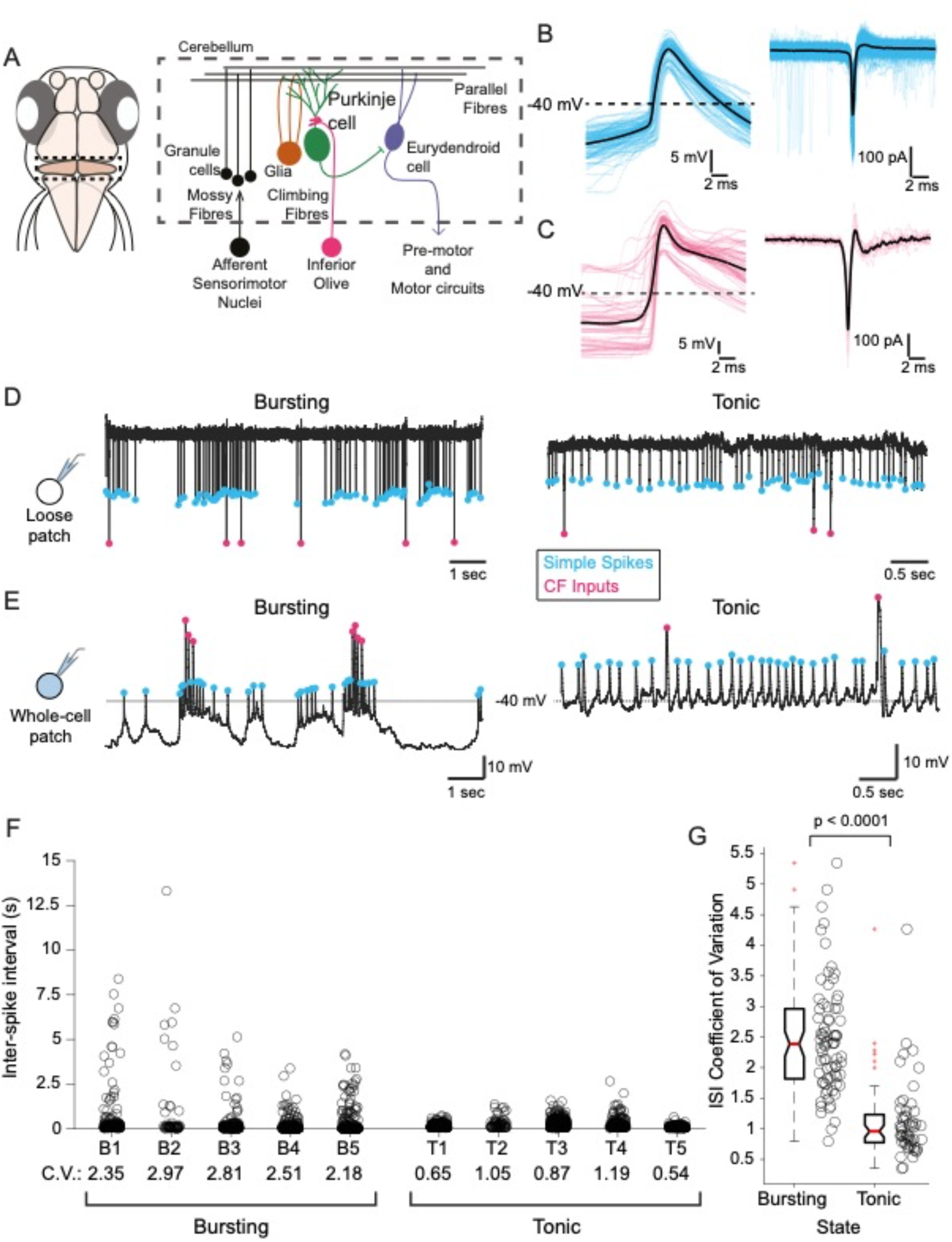
Cerebellar Purkinje neurons (PNs) in larval zebrafish exhibit membrane potential bistability (A) Schematic of the cerebellar circuitry in larval zebrafish (*Danio rerio*), showing the location of the cerebellum in the brain (left, dashed line box) and the cell types and circuit architecture (right). Purkinje neurons (green) are the principal neurons in this circuit and receive multiple inputs from afferent sensorimotor nuclei outside the cerebellum via parallel fibers (black) and climbing fibers (magenta). **(B)** Simple spikes, as observed intracellularly (left) and extracellularly (right). Individual events (n=100, intracellular and n=100, extracellular) are shown in cyan and have been aligned and superimposed on one another. The average of these events is shown in black. **(C)** Climbing fiber (CF) inputs, as observed intracellularly (left) and extracellularly (right). Individual events (n=55 intracellular and n=10 extracellular) are shown in magenta, and their respective averages in black. **(D)** Representative loose patch recordings from larval zebrafish PNs showing the two modes of firing. The bursting mode recording is shown on the left, and the tonic mode is shown on the right. In each mode, electrical events can be distinguished by their amplitude, with small-amplitude simple spikes marked in cyan and the large-amplitude CF inputs marked in magenta. **(E)** Representative whole-cell current clamp recordings showing the two modes of firing in PNs, with the same colour scheme for events as before. Cells shown in D and E are different. **(F)** Example plots of simple spike inter-event intervals from individual recordings. Five example cells are shown from each class (Bursting and Tonic). The coefficient of variation of this distribution is marked under each cell as “C.V.”. **(G)** The distribution of inter-spike interval coefficients of variation (n=131 cells), sorted by state (n=53 for tonic, and n=78 for bursting). The distributions are significantly different, tested using a Mann-Whitney U-Test (p = 6.0733e-16).

**Table 1.** Number of recordings compiled for the state-labelled imaging dataset.

|  | Extracellular | Intracellular | Total |
| --- | --- | --- | --- |
| Bursting | 39 | 39 | 78 |
| Tonic | 11 | 42 | 53 |
| Switch | 0 | 5 | 5 |
| Ambiguous | 0 | 25 | 25 |
| Poor Quality | 0 | 12 | 12 |
| Total | 50 | 123 | 173 |

Here, data was split into two groups - one that contained traces only with a single state throughout the recording, and another that contained traces with switches in state. Once split, the spike detection protocol described above was applied to each group of recordings to detect the simple spikes and CF inputs. These event timings were then used to generate simulated calcium imaging traces as described above. Combining the reconstruction with the assigned state resulted in the state-labelled calcium imaging dataset.

### Principal Components Analysis

Principal Components Analysis (PCA) was performed using the PCA module that is part of the *sklearn* package in Python (Pedregosa et al., 2011). The input to the model was a vector with the trace features for each 10-second-long trace in the reconstructions dataset (number of peaks, peak amplitude, area under the curve, mean, standard deviation and coefficient of variation). A 2-component PCA was performed, and the resulting fit (shown in Figure 4E) explained 88% of the variance in the data.

**Figure 2.**
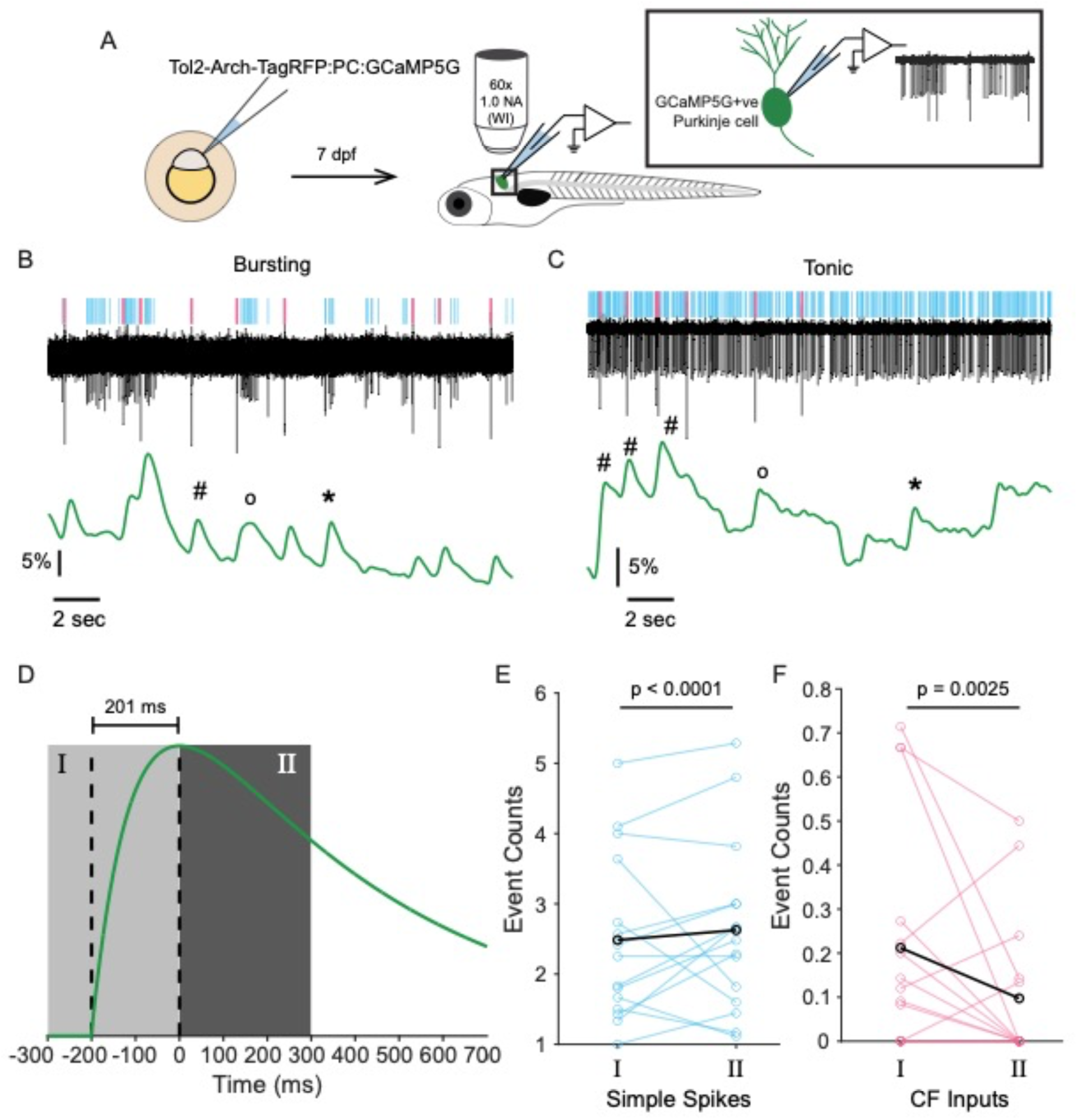
Both simple spikes and CF EPSPs contribute to the calcium signal in PNs, irrespective of cellular state. (A) Schematic of the experimental design and setup of simultaneous calcium imaging and electrophysiology in larval zebrafish. **(B)** and **(C)** Representative traces obtained during simultaneous imaging and electrophysiology for cells in the bursting (B) and tonic (C) mode. Simple spikes and CF inputs in the trace are identified and marked in the raster above the trace in cyan and magenta, respectively. The simultaneously-recorded change in GCaMP fluorescence (dF/F) is shown below in green. Calcium transients corresponding to simple spikes only (‘*’), CF-events only (‘#’), and both events (‘o’) are marked. **(D)** The typical GCaMP5G response to a single action potential, assuming it follows the profile of a difference of single exponentials with a half-rise time of 100ms and half-decay time of 500ms. The vertical dashed lines indicate the time of initiation of the calcium transient and the time when the transient peaks, which corresponds to an interval of 201ms. Region I is the period up to 300ms before the peak of the calcium signal, and region Ⅱ is the period up to 300ms after the peak of the transient. There were a total of n=171 transient peaks in the dF/F signal detected across the N=15 cells. **(E)** Average simple spike counts in the periods corresponding to regions Ⅰ and Ⅱ as defined in (D). Each cyan line corresponds to an individual cell (N=15) and the average of all these lines is shown in black. **(F)** Average CF input event counts in the periods corresponding to regions Ⅰ and Ⅱ as defined in (D). Each magenta line corresponds to an individual cell (N=15) and the average of all these lines is shown in black.

**Figure 3.**
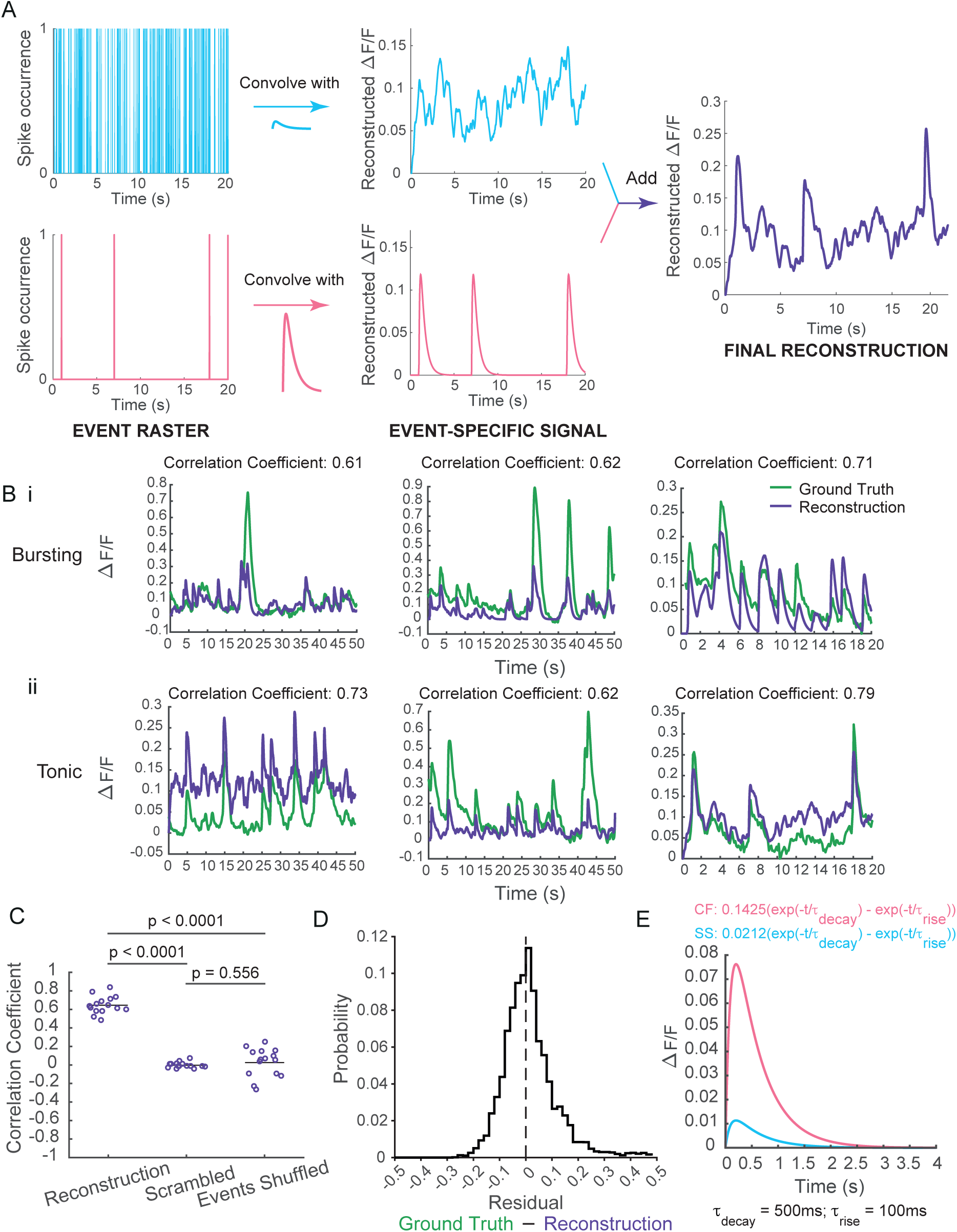
Reconstruction of the calcium signal from electrophysiology traces. (A) Schematic showing the method used to reconstruct the calcium signal from an electrophysiological trace. Simple spike (cyan) and CF input (magenta) rasters (left) are independently convolved with calcium sensor kernels (cyan for simple spikes, magenta for CF inputs) to generate an event-specific calcium signal time series (middle). These are then simply added together to get the final reconstruction (right). **(B)** Comparison of ground truth (green) and reconstructed (purple) calcium signals for 3 randomly chosen bursting cells (Bi) and 3 randomly chose tonic cells (Bii). The Pearson’s correlation coefficient between the two traces is marked above the plot for each pair. **(C)** Correlation coefficients for all cells calculated for either the true reconstruction, a scrambled version of the reconstruction, or the reconstruction for when simple spike and CF input event times were shuffled. A Kruskal-Wallis test yielded a p-value of 2.108 x 10^-7^, which was followed by a post-hoc Dunn’s test for individual comparisons, the latter of which is marked on the figure (n=15 cells; 6 bursting, 9 tonic). **(E)** Distribution of residuals between the ground truth and reconstructed calcium signal, pooled across all cells (n=15 cells). **(F)** The optimal calcium sensor kernels for simple spikes (cyan) and CF inputs (magenta). The formula for each kernel is mentioned above the plots, and the values for rise and decay time constants are mentioned below the plot.

**Figure 4.**
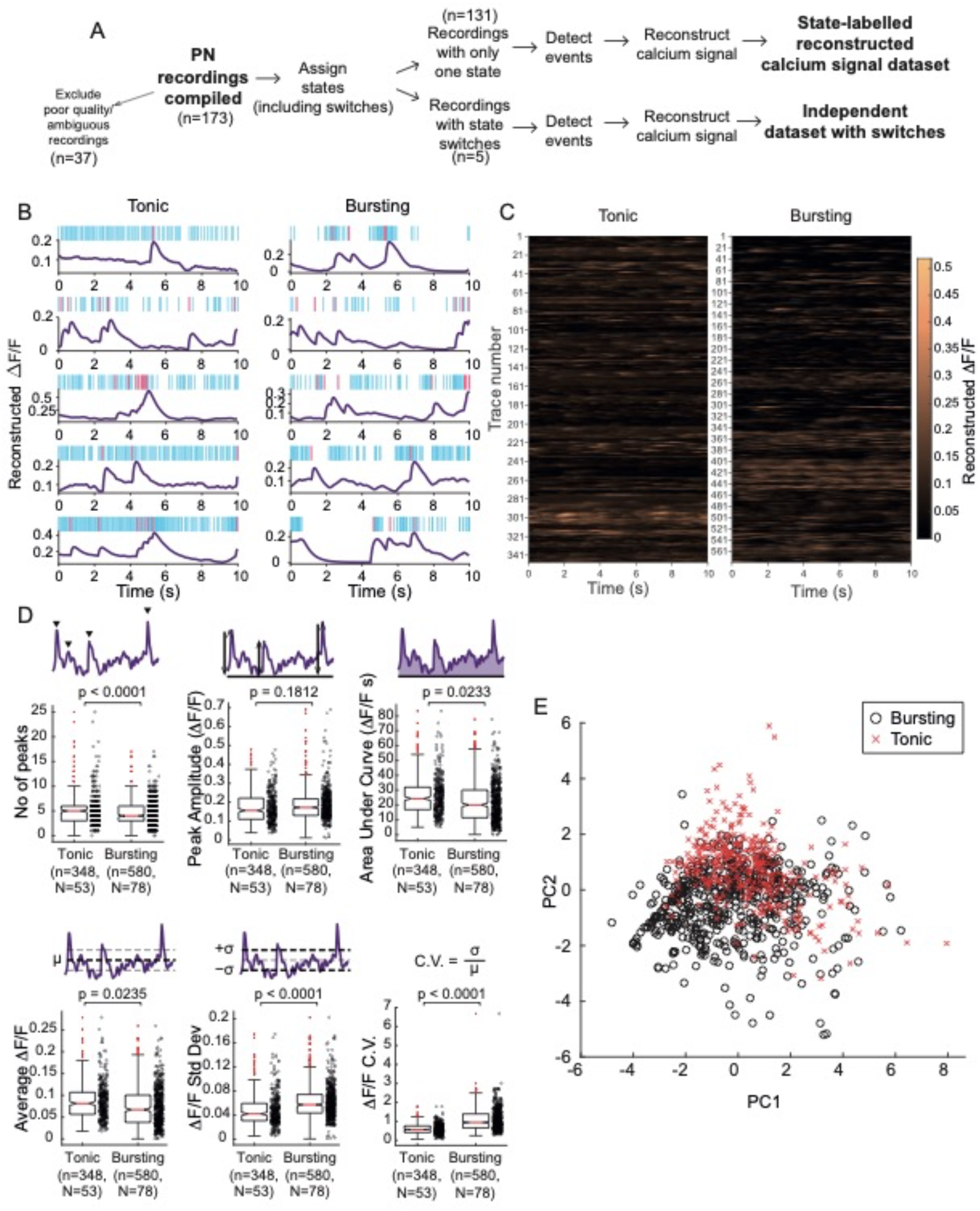
Generating a labelled dataset of reconstructed calcium signal traces. (A) Flowchart showing how electrophysiological recordings from PNs were processed to generate the state-labelled calcium signal database. **(B)** Representative randomly sampled traces from the state-labelled reconstructed calcium signal dataset. The left column has reconstructions from tonic cells and the right column has reconstructions from bursting cells. Above each trace is the raster showing the events in the source recording, with simple spikes in cyan, and CF inputs in magenta. **(C)** Heatmaps representing all the reconstructions in the state-labelled calcium signal dataset, for each state, after splitting all traces into 10-second-long non-overlapping chunks. (n=348 tonic traces; n=580 bursting traces). **(D)** Comparing the distribution of trace properties for both states. The top three plots show (from left to right) the number of peaks, peak amplitude and area under the curve of the reconstructions in the dataset. The bottom three plots show (from left to right) the mean, standard deviation and the coefficient of variation of ι1F/F values. n’s are the number of 10s-long samples. N’s are the number of cells used to generate the samples. p-values were computed using linear mixed-effects models. **(E)** Principal Components Analysis of trace properties, with data points marked by state - tonic (red crosses) and bursting (black open circles).

### Training the CNN-LSTM model for classification

The dataset of reconstructed calcium imaging time series was first split into 5 folds to perform cross validation (Fig. 5A). The experiments are repeated by considering each fold as a test set, with one of the folds being the validation set and remaining 3 folds to be training sets such that the data split for each experiment is as follows - 60% for training, 20% for validation, 20% for testing. This splitting was done at the level of entire traces, such that the training, testing and cross-validation sets all had a balanced representation of traces from both the tonic and bursting classes. We chose to do a 5-fold cross-validation, i.e. the entire training dataset is split into 5 parts, each containing 20% of the data. Four of the folds are used for training and the remaining one is used for cross-validation.

**Figure 5.**
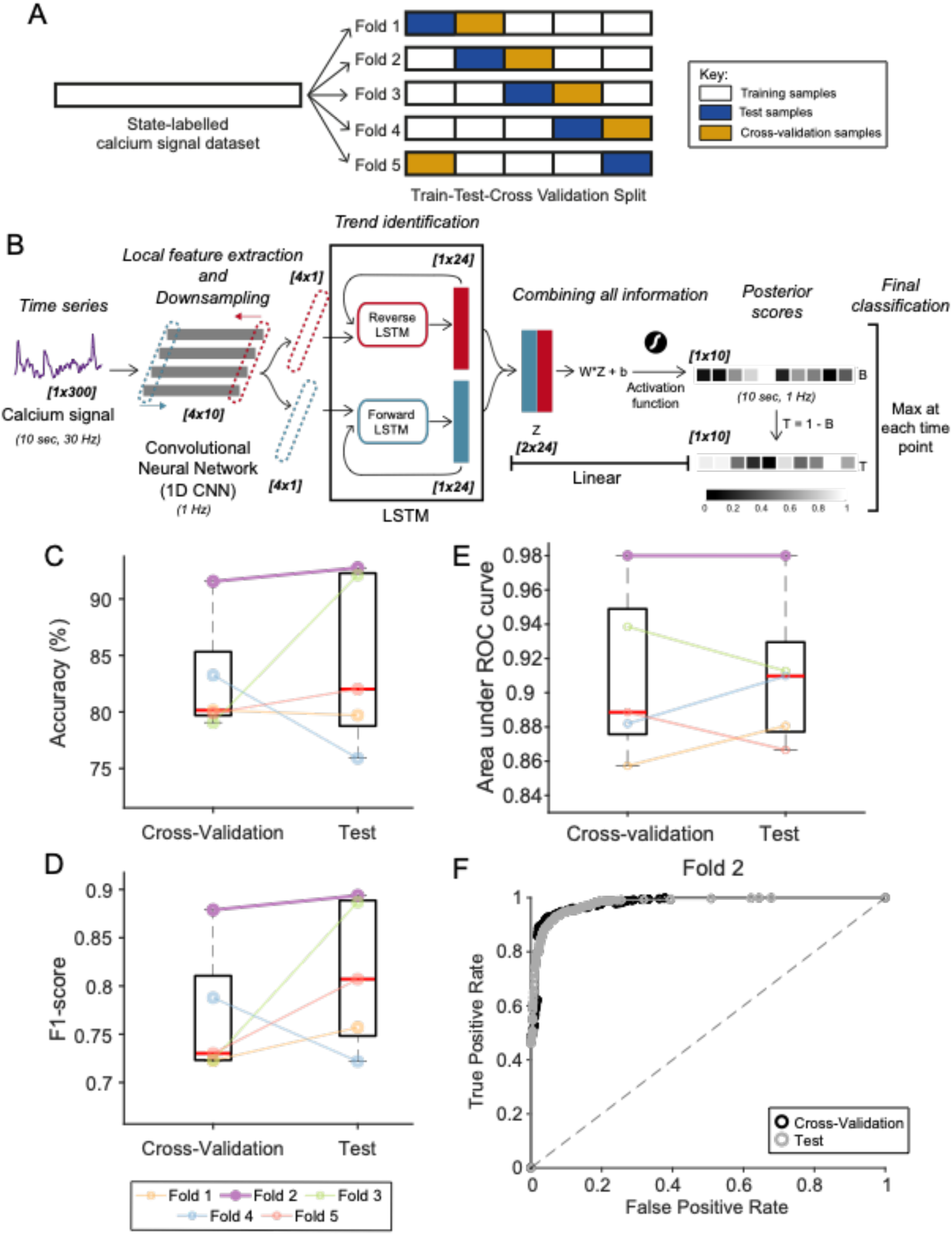
Design and training of CaMLsort (A) Flowchart showing how the state-labelled calcium signal dataset was split into training and test datasets using a 5-fold 60-20-20 split into training (white boxes), test (blue boxes) and cross-validation (yellow boxes) samples, respectively. The networks that performed well in the training phase, as assessed by performance on the cross-validation samples, were then challenged with the unseen test data (blue boxes) in each fold. An independent dataset with state switches was also used to further validate the networks obtained. **(B)** The architecture of the convolutional recurrent neural networks used to solve the classification problem. The network takes an input time series trace sampled at 30Hz (purple) and that is a multiple of 10 seconds long in duration (here just 10 seconds long). This trace is first normalized using min-max normalization, following which it is passed through a 1-dimensional convolutional neural network (1D CNN), which has 4 kernels with a step size and stride length of 30 each. The resulting 4 traces (grey boxes) are down-sampled, local feature-extracted versions of the original trace. All 4 outputs are passed to the LSTM module (central black box) for trend identification. The LSTM is a bidirectional one, which processes traces in both the forward (blue arrow) and reverse (red arrow) directions. A single time step is taken from all the CNN outputs (red and blue dotted boxes) and passed to the respective LSTM cell, which then processes the input to produce a resultant “hidden state” output (red and blue filled boxes). This output is passed back into the LSTM cell along with the next sample from the CNN’s output traces, until every time step from it has been processed (here 10 rounds). Once this is done, the final “hidden state” from each LSTM cell is retained and concatenated before their weighted average is taken by a Linear layer. The resulting average is converted to a “posterior score” using an activation function. These posterior scores represent the likelihood of the cell being in the bursting state at each time step. The likelihood of it being in the tonic state can be calculated from this. The final call is taken to be the state which has a higher posterior score. The text in italics represents the net effect of each phase of the neural network. The numbers in parentheses indicate the sampling frequency and/or the duration of the resultant vectors at various stages of processing by the neural network. Similarly, the numbers in bold and italics within square brackets indicate the size of the vectors/matrices at various stages of passing through the network. **(C)** Average classification accuracy of trained networks at the cross-validation and the test phases. Each fold is represented in a different color, marked in the legend at the bottom of panel (D). **(D)** Average F1-scores for each of the trained networks at the cross-validation and the test phases. **(E)** Area under the ROC (Receiver Operating Characteristic) curve for each of the five trained CNN-LSTM networks for both cross-validation (black) and test (gray) data. **(F)** ROC curve for Fold 2 of the CNN-LSTM for cross-validation (black) and test (gray) data.

The architecture of the network is shown in Fig. 5B. The input to the network is the entire trace sequence at 30Hz, normalized using min-max normalization for every non-overlapping 10 second period, thus setting the minimum to 0, and maximum to 1. Note that if the sampling frequency of a calcium trace was not 30Hz, it was first linearly interpolated to a 30Hz sampling frequency before being seen by the neural network. The first layer of processing is a CNN with kernel size 30 and stride size 30. The CNN down samples the input trace from 30Hz to 1Hz, and produces 4 vector outputs,which act as the input to the LSTM. The LSTM is bidirectional, so it looks at the sequence in both the forward and reverse order to produce an output. The LSTM has 24 hidden layers, so it produces 48 numbers per time step (Fig. 5B). The feature dimensions are collapsed by a Linear layer by taking their weighted average. This provides the classification logits at each position, which is passed through a softmax layer. The network is optimized using a cross entropy loss and the Adam optimizer (Kingma and Ba, 2017). The output of a trained neural network, then, is the probability of each timestep (at 1 Hz) belonging to the bursting class, from which the probability of it belonging to the tonic class can be derived. The class with the higher probability is assigned to each time step.

Since the model acts on the entire trace sequence, the amount of independent data for training the model is significantly reduced. To overcome this issue, we used a data augmentation strategy, which allows the network to learn features on arbitrary length sequences. For a sequence of length N, we determine a random segment size n, with n between [k, N], where k is the minimum sequence length. We randomly determine a starting position x, with a constraint that x < N - n. In our case, k=300 samples, and n varies in multiples of k (i.e. n = 300, 600, 900…). In each iteration of the network pass during the training phase, this random sequential segment was taken as the input trace to the model and gradient descent was performed to update weights. This was done for all the traces in the training set, so that the network saw entire traces at least once and in a non-overlapping fashion during the 500 epochs of training. The training is stopped before the full 500 epochs, based on early stopping criteria - if the validation loss does not improve after few epochs. All models were trained using the PyTorch framework in Python 3 (Paszke et al., 2019).

For all evaluations - cross-validation, test and other independent validations, the network had to make predictions for the entire trace duration and not just the n- sample long chunks. Networks were evaluated using two metrics - classification accuracies and F1-scores. Classification accuracies are simply the fraction of predictions that were correct. F1-scores, on the other hand, are adjusted for class imbalances and account for the kind of errors made. By definition, an F1-score is the harmonic mean of the precision and recall metrics, where precision is the fraction of all predicted positives that were true positives, and recall is the fraction of all ground truth positives that were true positives.

To improve/smoothen predictions made by CaMLsort, we used a majority voting strategy. To do this, we re-sampled the time series using rolling window sampling with a window length *N*. For each time step of prediction, *t_n_*, we found all the *m* windows (where *m*≤*N*) that contained *t_n_* (i.e. [*t_n-N+1_*,*t_n-N+2_*,… *t_n_*,*t_n+1_*], [*t_n-N_*,*t_n-N+1_*,… *t_n-1_*,*t_n_*], and so on until [*t_n_*, *t_n1_*,… *t_n+N-1_*,*t_n+N_*]). For each of these windows, an average posterior score was calculated for each state, and the “average class” taken to be the state which had the higher score. The “majority class”, then, was the one which was the most frequent average class across all *m* windows. A “confident call” was made if and only if the average posterior scores for the majority class across all *m* windows was strictly higher than the average posterior scores for the minority class.

### Generating artificial Purkinje neuron spike trains

Synthetic spike trains of the tonic class were generated using a Poisson statistical model. The only parameter needed for this is an average event rate, λ, which we took to be a random number between 4-10 for simple spikes. Spike trains for bursting neurons do not follow Poisson statistics. Hence, we used an inverse sampling method to generate spike trains of the bursting class. For this, we used the compiled electrophysiology data to obtain the real cumulative distribution of interevent intervals for simple spikes across all bursting cells. A random number, y, is generated from the uniform distribution and it is used to find a value x from the reference cumulative interspike interval (ISI) distribution such that P(ISI <= x) = y.

Thus, by generating events sequentially with a given next interevent interval, we obtained artificial simple spike trains for the bursting class.

CF input event trains were generated using a Poisson statistics model with an average event rate between 0.2-1. The distributions of interevent intervals in the synthetic datasets were compared against the real distributions using histograms and QQ plots. Independent reviewers were asked to classify unlabelled spike trains as real or artificial and managed to do so with an accuracy of ∼42% on average.

### Statistical Analysis

We used a one-sided Mann-Whitney U test to compare the distributions of interspike interval coefficient of variation between tonic and bursting cells, for both Purkinje neurons (Fig 1G) and dopaminergic VTA neurons (Fig. 9C).

**Figure 6.**
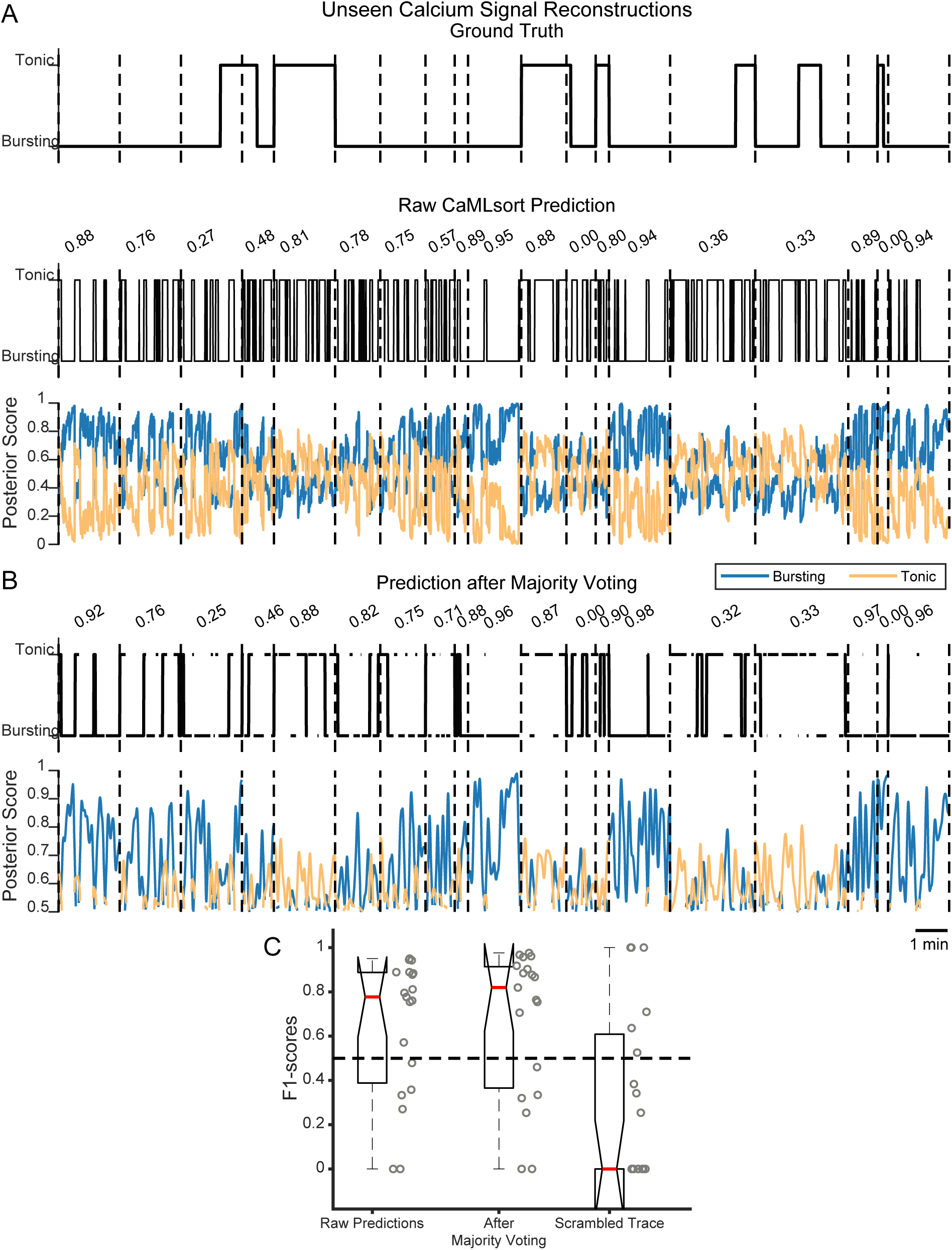
CaMLsort predictions from previously unseen calcium signal reconstructions (A) Raw CaMLsort class label predictions from previously unseen calcium signal reconstructions (n=19 recordings). Predictions from CaMLsort (middle) are compared against the ground truth state (top) at each time step. Predictions for each recording were made independently, but have been stitched here for the purposes of representation. Each recording is separated by a vertical dashed line. The number above each prediction is the F1-score for that recording. The bottommost plot has posterior scores for each time step, with those for the “bursting” class in blue, and those for the “tonic” class in light orange. **(B)** CaMLsort predictions (top) that remain after majority voting with a window length of 7 steps has been used to retain only confident classifications (see Results and Methods for details). The posterior score (bottom) for each class at each time step, taken as the average class posterior score across all 7-step windows that included said time step. As in (A), the posterior scores for the “bursting” class are in blue, whereas those for the “tonic” class are in light orange, and the numbers above each prediction represent the F1-scores for that recording. **(C)** The distribution of F1-scores between the ground truth state and various predictions from CaMLsort - either the raw predictions (left), predictions obtained after majority voting (middle), or the raw predictions obtained by passing a scrambled version of the calcium signal as an input.

**Figure 7.**
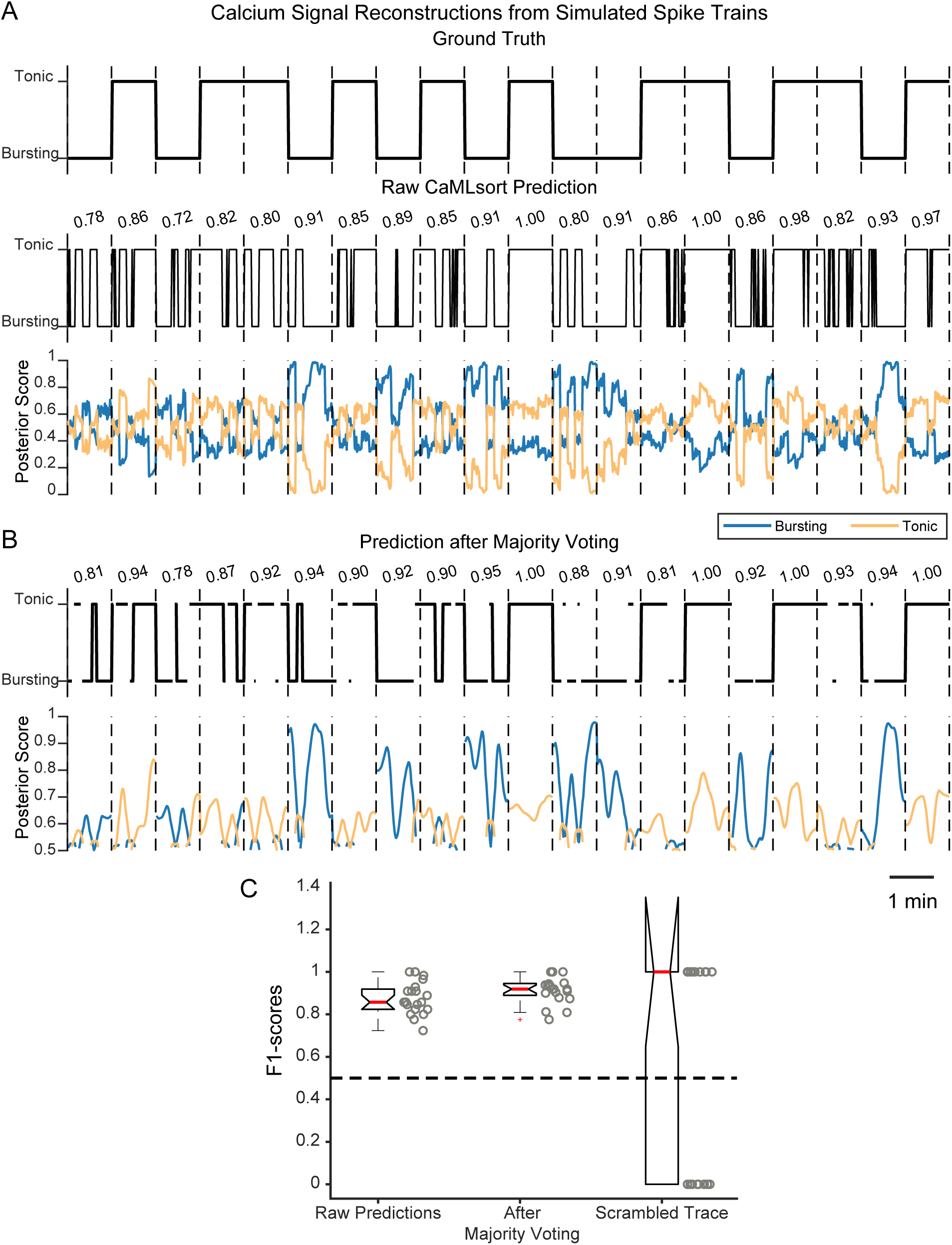
CaMLsort predictions from calcium signal reconstructions of simulated spike trains (A) Raw CaMLsort class label predictions from previously unseen calcium signal reconstructions that were generated from simulated spike trains (n=20 recordings). Predictions from CaMLsort (middle) are compared against the ground truth state (top) at each time step. Predictions for each recording were made independently, but have been stitched here for the purposes of representation. Each recording is separated by a vertical dashed line. The number above each prediction is the F1-score for that recording. The bottommost plot has posterior scores for each time step, with those for the “bursting” class in blue, and those for the “tonic” class in light orange. **(B)** CaMLsort predictions (top) that remain after majority voting with a window length of 7 steps has been used to retain only confident classifications (see Results and Methods for details). The posterior score (bottom) for each class at each time step, taken as the average class posterior score across all 7- step windows that included said time step. As in (A), the posterior scores for the “bursting” class are in blue, whereas those for the “tonic” class are in light orange, and the numbers above each prediction represent the F1-scores for that recording. **(C)** The distribution of F1-scores between the ground truth state and various predictions from CaMLsort - either the raw predictions (left), predictions obtained after majority voting (middle), or the raw predictions obtained by passing a scrambled version of the calcium signal as an input.

**Figure 8.**
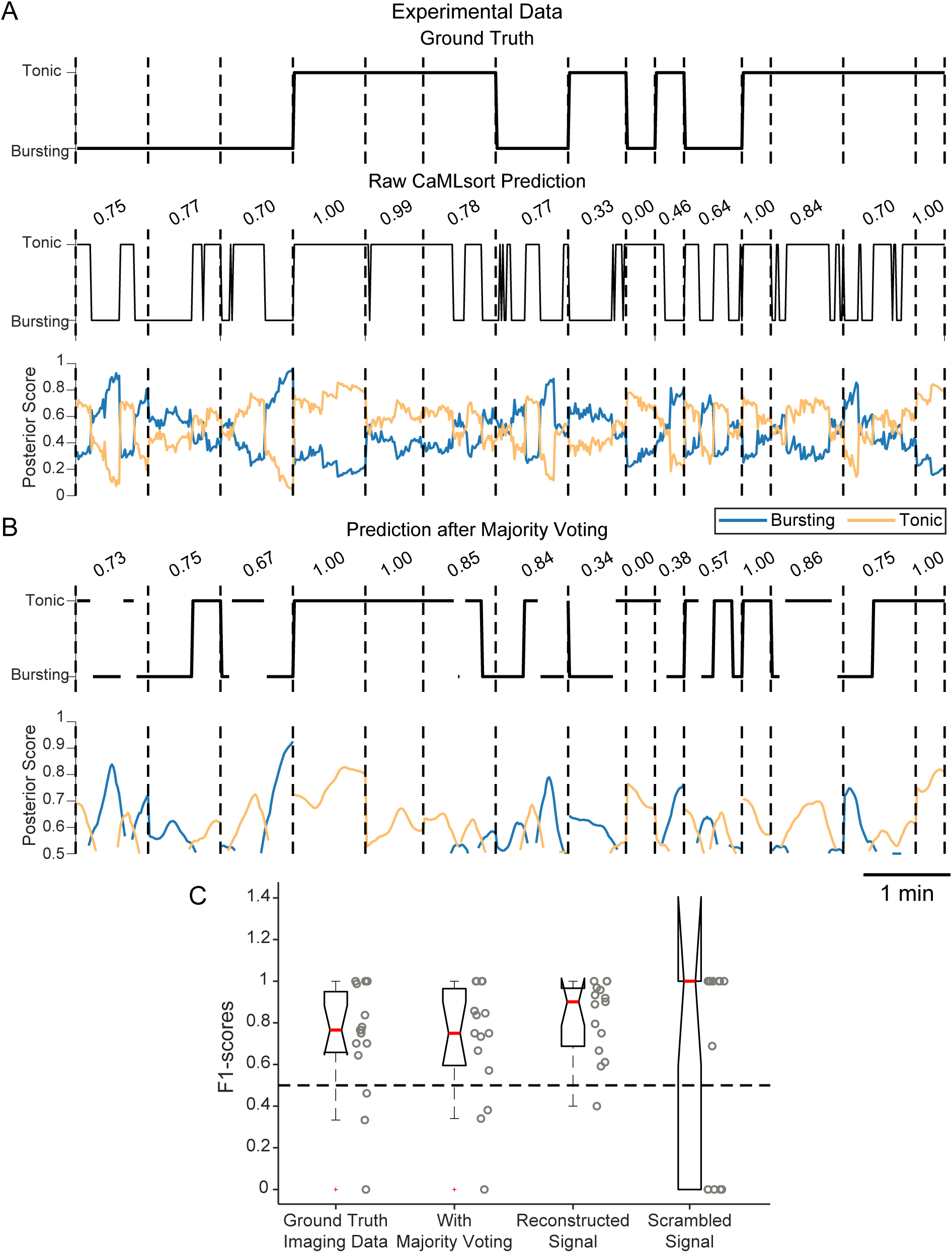
CaMLsort predictions on imaging data from zebrafish Purkinje neurons (A) Raw CaMLsort predictions of cellular state inferred from experimental data acquired using widefield imaging (n=15; also see Chapter 4). Predictions from CaMLsort (middle) are compared against the ground truth state (top) at each time step. Each recording was treated independently, but predictions across cells have been stitched here for ease of visualisation. Each recording is separated by a vertical dashed line, and the number above each prediction is the F1-score for that recording. The bottommost plot has posterior scores for every time step, with those for the “bursting” class in blue, and those for the “tonic” class in light orange. **(B)** CaMLsort predictions (top) that remain after majority voting with a window length of 7 steps has been used to retain only confident classifications (see Results and Methods for details). The posterior score (bottom) for each class at each time step, taken as the average class posterior score across all 7-step windows that included said time step. As in (A), the posterior scores for the “bursting” class are in blue, whereas those for the “tonic” class are in light orange. **(C)** The distribution of F1-scores between the ground truth state and various predictions from CaMLsort - raw predictions from the imaging data (far left), predictions obtained after majority voting (second from left), raw predictions obtained from calcium signals reconstructed from spike times (third from left) and those predictions from imaging dF/F traces that were scrambled prior to prediction.

**Figure 9.**
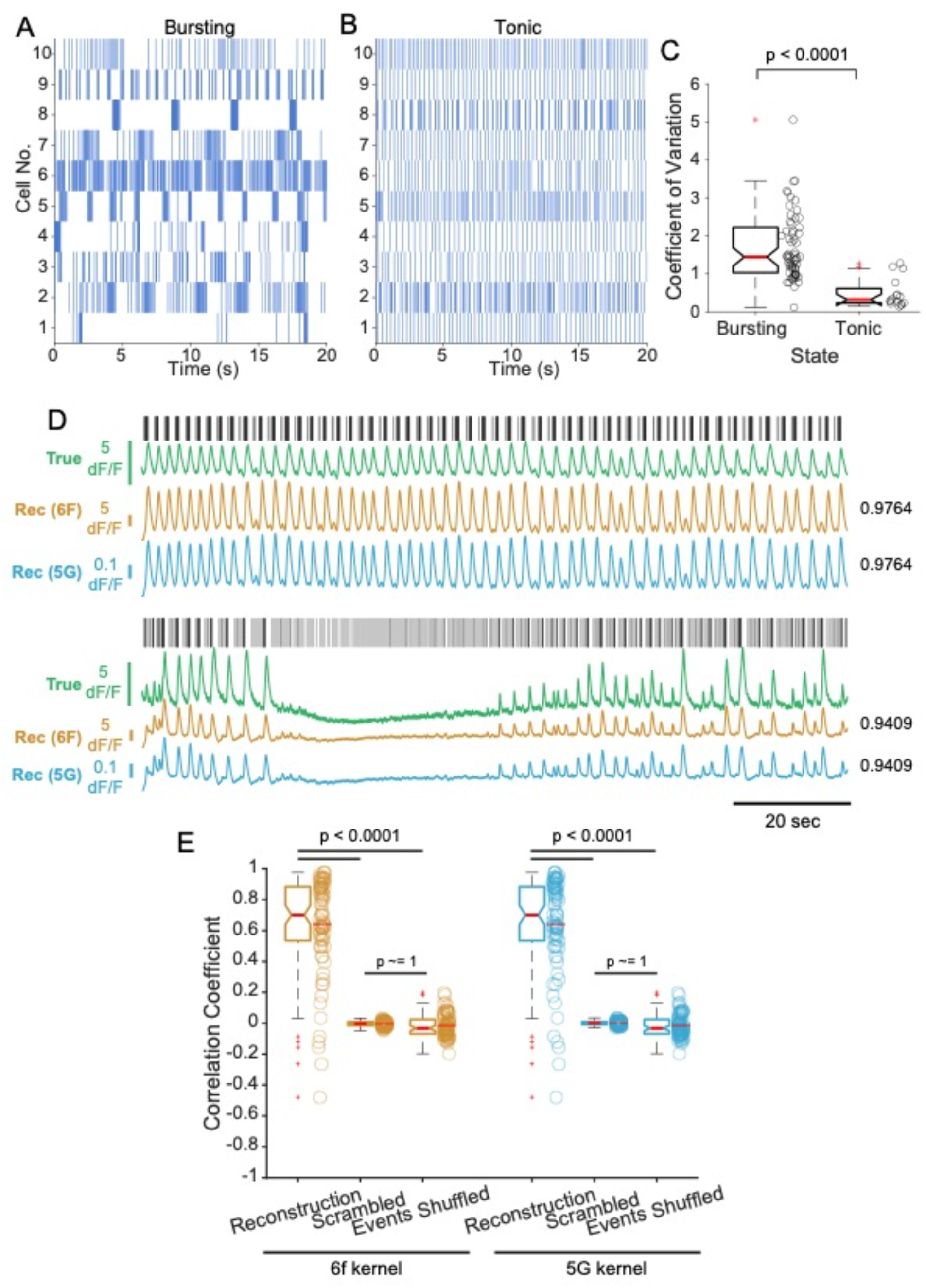
Reconstructing calcium signals for VTA DA neurons using a convolution-based forward model. (A-B) Spike rasters of two representative VTA DA neurons. **(C)** The distributions of C.V.s of interspike intervals for cells classified as either bursting (n=62) or tonic (n=16 cells). The p-value was calculated using a one-sided Mann-Whitney U-test. **(D)** Electrophysiological activity and associated calcium signals from two representative VTA DA neurons. For each cell, spike timings are shown as rasters (black) and the raw simultaneously-acquired dF/F signals aligned below (green). Two calcium signal reconstructions were generated, using either a GCaMP6f kernel (gold) or a GCaMP5G kernel (blue). Numbers alongside each reconstruction indicate the Pearson correlation coefficient between the ground truth and the corresponding calcium signal reconstruction. **(E)** Quantification of the reconstruction quality, as measured using the Pearson correlation coefficient between the ground truth and reconstructed signals (“Reconstruction”). For each cell (n=78), the correlation between ground truth and a scrambled version of the reconstructed signal (“Scrambled”), as well as the correlation between ground truth and a reconstruction generated after shuffling spike timings (“Events Shuffled”) were measured. Individual data points are plotted alongside each boxplot. p-values were calculated using a Kruskal-Wallis test yielded a p-value of 4.6130 x 10^-49^, and then using the post-hoc Dunn’s test for individual comparisons, the latter of which are marked on the figure.

To compare the quality of reconstructions obtained from the generative forward model, we used a Kruskal-Wallis test to detect any overall group differences and followed it up with a post-hoc Dunn’s test for individual comparisons (Fig. 3C and Fig. 9E).

For comparisons involving a nested data structure, such as those involving trace features between tonic and bursting cells (Fig. 4D), where there are multiple cells and multiple measurements per cell, we used a linear mixed-effects model with a maximum likelihood estimator to calculate p-values.

## Results

### Zebrafish Purkinje neurons exhibit membrane potential bistability

Purkinje neurons (PNs) are the principal neurons in the cerebellar circuit (Fig. 1A). In larval zebrafish, they exhibit two kinds of electrical events that can be distinguished solely based on their amplitude in both extracellular and intracellular recordings (Sengupta and Thirumalai, 2015) (Fig. 1 B-E). The smaller amplitude events are action potentials and are called simple spikes (Fig. 1B,D,E, cyan), while the larger amplitude events are all-or-none synaptic input from climbing fibers (Fig. 1C,D,E, magenta). Individual simple spike and CF input waveforms are shown in Fig. 1B and 1C, respectively, for both whole-cell and juxtacellular recordings.

These neurons exhibit membrane potential bistability, i.e. they are found to be at one of two stable membrane potentials, depending on which they fire either tonically or in bursts. Cells can spontaneously switch between the two states, but can also be taken from one state to the other by current injection (Sengupta and Thirumalai, 2015).

The two states can be distinguished visually from extracellular loose-patch recordings based on (i) the firing frequency of simple spikes and (ii) the presence of long intervals of inactivity in the burst mode (Fig. 1D, left) which are not present in the tonic mode (Fig. 1D, right). The same features can be used to distinguish between the two modes in whole-cell patch- clamp recordings, along with an extra feature – membrane potentials are more hyperpolarized in the bursting mode (Fig. 1E, left) as opposed to the tonic mode (Fig. 1E, right). Additionally, in tonic cells, inter-spike intervals form a single cluster, while bursting cells have inter-event intervals that trail to large values, corresponding to the inter-burst intervals (Fig. 1F). This is also reflected in the values of coefficient of variation of the inter-spike intervals, which are larger in bursting cells than the more regularly spiking tonic cells (Fig. 1G). Given that there is a lot of variability across cells and recordings, no single cut-off value could be determined for these parameters (Fig. 1G). Instead, all the aforementioned features were taken together to determine the final state classification.

### Calcium imaging reports both simple spike bursts and individual CF inputs, independent of state

Given that PNs manifest two distinct activity patterns (tonic and bursting) and two types of electrical events (simple spikes and CF events), we wondered if calcium imaging signals can reliably report these distinct patterns of activity. To do this, we performed simultaneous wide-field calcium imaging and targeted patch-clamp electrophysiology from zebrafish PNs. We used the carbonic anhydrase (Ca8) enhancer element and a minimal promoter (*cfos*) to drive expression of the genetically encoded calcium sensor GCaMP5G in PNs (Matsui et al., 2014).

Embryos were microinjected at the 1-2 cell stage and those larvae that showed sparse labelling in PNs were selected for targeted patch-clamp electrophysiology and imaging (Fig. 2A). We chose to perform extracellular, loose-patch recordings so as to be minimally invasive and to avoid the dialysis of cellular contents including GCaMP. We obtained such recordings from 15 cells in 12 larvae, out of which 9 cells were naturally in the tonic state, and 6 cells in the bursting state.

Representative traces from this experiment for cells in the bursting and the tonic states are shown in Fig. 2B and 2C, respectively. The first observation we made was that many of the calcium transients (marked with a hash symbol in Fig. 2B and 2C) had an almost one-to-one correspondence with CF inputs (magenta lines in the raster plot) in cells of both states. In tonic cells, most simple spikes were not reported as transients, as one would expect from cells that fire at high frequencies. However, in some cases, calcium transient peaks occurred in the absence of CF inputs, during times of intense simple spike firing (Fig. 2C, see asterisk and cyan lines in raster plot). Similarly, in bursting cells, we found some calcium transients associated only with a burst of simple spikes (Fig. 2B, asterisk). Naturally, then, one would expect to see calcium transients when CF inputs co-occurred with a burst of simple spikes, which is what we observed (marked with open circles in Fig. 2B and 2C). Thus, regardless of the state of a cell, we found calcium transients that corresponded to individual CF inputs, bursts of simple spikes or a combination of both.

While this method confirms that a calcium transient occurs after a CF input, suggesting a causal link between the CF input’s arrival and the measured calcium influx into the cell, it is not useful in the case of simple spikes. Firstly, there are multiple simple spikes per frame - approximately 2-3 spikes per frame when imaged at 30Hz. This is further worsened by the timescales of GCaMP5G’s rise and decay, which occurs on the order of 100s of milliseconds (Akerboom et al., 2012; Chen et al., 2013). Moreover, one cannot separate out the individual effects of simple spikes and CF inputs because the two events are interspersed and thus, when calculating the triggered average response for one event, there is bleed-through from the other.

To ascertain whether CFs and simple spikes indeed trigger calcium signals, we decided to compare the number of events in the time leading up to the calcium peak (Fig. 2D, region Ⅰ) and during the decay period (Fig. 2D, region Ⅱ). The expectation was that if an event triggered calcium activity, there would be more of them in region Ⅰ, which includes the time of initiation of the calcium transient (Fig. 2D), than in region Ⅱ, when the signal is decaying. GCaMP5G has a rise time constant of ∼100ms, which means that the time it takes to reach a peak value from the baseline is ∼201ms (modelled using a difference of single exponentials, see Methods).

Hence, to be sure that the time of initiation was included in region Ⅰ, we considered a 300ms interval before the peak of the calcium signal. An interval of the same 300ms duration after the peak was taken to be region Ⅱ.

We first detected calcium transient peaks in the dF/F signal (see Methods), of which there were a total of n=171 detected across the N=15 cells. The average number of simple spikes (Fig. 2E) and CF inputs (Fig. 2F) are significantly different in region I and II (linear mixed effect model; p<0.0001 for simple spikes and p = 0.0025 for CF inputs). There are more CF inputs in region I compared to region II indicating that these events can trigger calcium signals. However, for simple spikes, there is a small, but significant increase in region II compared to region I. While this is opposite of what one might have expected, it can be explained by the high rate of simple spikes lasting over several hundreds of milliseconds in both bursting and tonic cells. In both regions I and II, there are approximately 2.5 simple spikes within the 300 ms window. These results imply that both simple spikes and CF inputs are associated with dF/F transients.

In summary, it is difficult to infer the underlying event train or even the cellular state of a Purkinje neuron directly from its calcium signal, as there is no one-to-one correspondence between calcium transients and their underlying electrophysiological events, as has also been reported previously (Knogler et al., 2019; Sengupta, 2015).

### The calcium signal for a Purkinje neuron can be reconstructed from its electrophysiological recording

Though the calcium signal in PNs could arise from more than one electrical event, we hypothesized that there might be sufficient information in the calcium signal time series to allow us to determine whether it is tonically firing or bursting. One possible approach to this problem could be the application of machine learning algorithms to build a classifier that labels cells as tonic or bursting given a time series of calcium transients. For this, it was first necessary to generate a large ground truth dataset (100’s of cells) comprising the electrical activity patterns and their corresponding calcium transients for training the classifier. Ideally, this dataset would take the form of what is shown in Fig. 2B,C, where both electrophysiology and imaging data are simultaneously acquired. Our ground truth dataset has only 15 cells, however, given that the simultaneous imaging and electrophysiology experiments are low yield and technically challenging. Hence, we needed a strategy to augment our dataset.

Mathematically, the calcium signal obtained by imaging can be thought of as the result of the convolution of spike trains with a calcium sensor kernel having a fast rise and a slow decay and described as a difference of exponentials. This transformation has been shown to work well in cells that are mostly quiescent and where every single action potential produces a large increase in intracellular calcium concentration (Smetters et al., 1999; Rupprecht et al., 2021; Vogelstein et al., 2010; Yaksi and Friedrich, 2006). We hypothesized that a similar transformation could be used to reconstruct the calcium signal profiles of PNs from their electrophysiological recordings alone. Such reconstructed calcium imaging traces with their corresponding electrical recording traces may then be applied to build the classifier.

However, given that these neurons have not one, but two electrophysiological events, each contributing differently to the calcium signal obtained from imaging, we must assign each event a separate kernel (Fig. 3A), which we assume to be independent of each other. This is a valid assumption, since simple spikes and CF events arise from distinct intrinsic mechanisms (Sengupta and Thirumalai, 2015). We used simultaneous imaging and electrophysiology recordings to find optimal kernels for simple spikes and CF events. For each PN recording, we obtained simple spike and CF event rasters. Each event raster was then independently convolved with its own exponential response kernel, and the resulting “event-specific signals” were linearly summed to produce the final reconstruction (Fig. 3A). We then calculated the correlation coefficient and the mean squared error (MSE) between the reconstructed calcium signal time series and the experimentally obtained calcium imaging time series.

With this description in place, the problem then becomes one of finding optimal coefficients for each response kernel. This optimization was done by gradient descent to minimize the MSE between the ground truth and the reconstruction (Fig. 3A). The process was performed independently for each cell in the dataset. Then, these optimal kernel combinations were used to make reconstructions for other cells in the dataset, and the kernel that generalized best across all cells, as quantified by least total MSE and highest total cross-correlation between the ground truth and reconstruction, was taken to be the best kernel for the nextsteps.

Although this is a simple forward model with calcium dynamics assumed to be linear and time-invariant, we found that this method to approximate the calcium signal works well, no matter the state of the neuron (Fig. 3B). The quality of the reconstructions was quantified by measuring the correlation coefficient between the ground truth and the reconstruction (Fig. 3C). To benchmark the correlation coefficient values obtained against those obtained simply by chance, we also calculated the correlation coefficient between the ground truth and a scrambled version of the reconstruction, as well as that between the ground truth and a reconstruction generated after shuffling all the event times in the recording (Fig. 3C). In both the latter cases, the correlation disappears, suggesting that the reconstructions generated by this forward model are, indeed, faithful reconstructions. We next calculated the residuals between ground truth and reconstructions, the distribution for which is shown in Fig. 3D. It is not normally distributed, and has a positive skew, which means that our reconstructions tend to underestimate the true signal. Nevertheless, the small values of the residuals further lend support to the idea that the forward model of calcium signal reconstruction works well.

Given that only groups/bursts of simple spikes produce a calcium transient amplitude comparable to that generated by an individual CF input (Knogler et al., 2019; Fig. 2B), one would expect a much smaller response kernel for simple spikes than for the CF input. Indeed, the optimal kernels found by this method show that the CF input response kernel is approximately 7.5 times larger than the simple spike response kernel (Fig. 3E).

### A state-labelled calcium signal dataset was generated by reconstructing the calcium signal for a compilation of electrophysiological traces

Having a method of converting any given PN electrophysiology trace to its corresponding calcium imaging trace allowed us to generate simulated calcium imaging traces for a compilation of all electrophysiological recordings acquired in our lab (Fig. 4A). To do so, we first pooled all the electrophysiological recordings, both extracellular and whole-cell, made during the period of 2011-2020, for a total of N=173 recordings. Recordings of poor quality (N=37), i.e., those with interruptions, or noisy baselines, unusually broad/filtered events, or indistinguishable simple spikes and CF inputs were first excluded from the dataset. Retained recordings then had a state assigned to them (using criteria described above; also see Fig 1D-G), including state switches, if there were any. This collection was split into two sets, one with those recordings which had only one state throughout (N=131), and another with those which had a state switch (N=5). The entire complement of recordings and their corresponding states is summarized in Table 1. Events detected in all the recordings were used to reconstruct a putative calcium signal using the kernels described in Fig. 3E. This finally produced two labelled datasets - a state-labelled reconstructed calcium signal dataset and an independent dataset with switches (Fig. 4A). The recordings used to generate this dataset were of different durations with a median duration of 60 seconds. In order to get multiple samples per recording, we decided to split the traces into 10- second-long non-overlapping chunks, representative samples from which are shown in Fig 4B. This subsampling yielded a dataset of state-labelled calcium imaging traces (n=928 traces of which n=580 bursting, n=348 tonic) that is much larger than could have been obtained experimentally. The entire complement of non-overlapping samples is shown as a heatmap in Fig. 4C.

To test whether there were properties of these traces that could be used to unambiguously classify their source state, we compared the number of peaks, average peak amplitude, area under the curve, mean, standard deviation and coefficient of variation of the reconstructed calcium traces (Fig. 4D). Except peak amplitude, all other properties were significantly different from each other for the tonic and bursting groups. Nevertheless, given any of these properties, it was not possible to unambiguously assign state as the distributions are overlapping (Fig. 4D). Further, a principal component analysis of these trace features also failed to separate tonic and bursting cells based on the above-mentioned features of their calcium traces (Fig. 4E).

Machine learning methods have successfully been used to solve such problems when the data points are not linearly separable (He et al., 2016). Thus, we turned to developing a machine learning model to identify the state of a cell from a calcium signal time series.

### Training a convolutional recurrent neural network to distinguish state from calcium imaging data alone

To find good machine learning models to solve this classification problem, we performed 5-fold cross validation, which is used to ensure that models generalize well and do not overfit the training data (Fig. 5A). On each fold, only 60% of the training traces were shown to the network in the training phase (Fig. 5A, white boxes), the network optimized on 20% of data used as a validation set (Fig. 5A, yellow boxes), and the generalizability of the trained network assessed using the remaining 20% (Fig. 5A, blue boxes). Each of the 5 folds had a different 60-20-20 split, such that the training and cross-validation datasets had a balance of traces from both classes.

We tested several different machine learning models on this classification problem - Support Vector Machines (SVMs), Deep Neural Networks (DNN), Convolutional Neural Networks (CNNs) and convolutional recurrent neural networks (CNN-LSTM, i.e. a Convolutional Neural Network with Long Short-Term Memory). The prediction accuracies and F1 scores (a metric that adjusts for class imbalances as well as whether a prediction was a true/false positive/negative, defined in the methods section) for the various models tested are shown in Table 2. Of these models, the 1D-CNN and CNN-LSTM models performed best across all folds. However, the former only predicts from snapshots of 10s-long snippets of the calcium trace, and the networks had more erroneous performance when predicting on the entire time series (data not shown). The CNN-LSTM model, however, also incorporates a recurrent neural network module (Hochreiter and Schmidhuber, 1997), which has the ability to learn features across an entire sequence and not just across neighbouring positions in a trace sequence, like CNNs do.

**Table 2.** Prediction accuracies and F1-scores for various models trained tosolve the classification problem.

| Model | Fold | Validation |  | Test |  |
| --- | --- | --- | --- | --- | --- |
|  |  | Accuracy (%) | F1-score | Accuracy(%) | F1-score |
| Support Vector Machine (SVM) | 1 | 81.41 | 0.7206 | 70.19 | 0.6077 |
|  | 2 | 72.68 | 0.593 | 79.66 | 0.6842 |
|  | 3 | 75.84 | 0.703 | 72.55 | 0.6055 |
|  | 4 | 73.10 | 0.6422 | 73.65 | 0.671 |
|  | 5 | 70.67 | 0.629 | 74.89 | 0.6755 |
|  | Mean ± Stdev | 74.74 ± 4.16 | 0.6576 ± 0.0531 | 74.19 ± 3.51 | 0.6488 ± 0.0388 |
| Deep Neural Network | 1 | 83.91 | 0.7893 | 72.99 | 0.6649 |
|  | 2 | 77.71 | 0.7205 | 82.89 | 0.7849 |
|  | 3 | 74.85 | 0.7099 | 76.10 | 0.6788 |
|  | 4 | 73.31 | 0.6535 | 75.12 | 0.7143 |
|  | 5 | 77.82 | 0.7060 | 75.24 | 0.6629 |
|  | Mean ± Stdev | 77.52 ± 4.06 | 0.7158 ± 0.0486 | 76.47 ± 3.77 | 0.7012 ± 0.0511 |
| 1D CNN | 1 | 87.20 | 0.8163 | 84.00 | 0.7933 |
|  | 2 | 80.67 | 0.7541 | 88.33 | 0.8336 |
|  | 3 | 82.66 | 0.7880 | 80.09 | 0.7263 |
|  | 4 | 78.28 | 0.7185 | 81.07 | 0.7719 |
|  | 5 | 82.60 | 0.7604 | 78.34 | 0.7004 |
| | Mean ±<br>Stdev | 82.28 ±<br>3.28 | 0.7675 ±<br>0.0369 | 82.37 ± 3.91 | 0.7651 ±<br>0.0530 |
| 1D CNN (3x) | 1 | 87.54 | 0.8226 | 84.00 | 0.7893 |
|  | 2 | 80.61 | 0.7535 | 86.97 | 0.8142 |
|  | 3 | 80.61 | 0.7650 | 78.41 | 0.7089 |
|  | 4 | 78.62 | 0.73 | 73.66 | 0.6592 |
|  | 5 | 82.49 | 0.7592 | 76.83 | 0.6775 |
| | Mean ±<br>Stdev | 81.97 ±<br>3.40 | 0.7661 ±<br>0.0343 | 79.97 ± 5.42 | 0.7298 ±<br>0.0686 |
| CNN-LSTM | 1 | 80.15 | 0.7227 | 79.7 | 0.7570 |
|  | **2** | **91.55** | **0.8788** | **92.75** | **0.8937** |
|  | 3 | 79.06 | 0.7233 | 92.12 | 0.887 |
|  | 4 | 83.25 | 0.7878 | 75.9 | 0.7219 |
|  | 5 | 79.9 | 0.7302 | 82.02 | 0.8071 |
| | Mean ±<br>Stdev | 82.78 ±<br>5.15 | 0.7686 ±<br>0.0674 | 84.50 ± 7.57 | 0.8133 ±<br>0.0766 |

The architecture of the network is shown in Fig. 5B. The input to the network is a trace sampled at 30Hz, which is some multiple of 10 seconds in duration. It is first remapped to values between 0 and 1 using min-max normalization and then processed by a CNN, which acts as a local feature extractor that also down samples the input data, such that it produces 4 vector outputs at a 1Hz sampling frequency. These outputs are passed to a bidirectional LSTM, one time-step at a time. A forward LSTM cell looks at trends in the forward direction (Fig. 5B, blue dotted box and arrow), while a reverse LSTM cell goes through the data in the reverse direction (Fig. 5B, red dotted box and arrow). Each of the two cells produces a “hidden state” as its output. Given that the LSTM is a recurrent neural network, this output is fed back into the network along with the next appropriate time step from the CNN’s output, a process which is repeated 9 more times (for every 10 second trace). In essence, the LSTM is detecting trends across the entire sequence. The final hidden states from the forward and reverse LSTMs, produced after passing through the entire trace, are retained and concatenated. These outputs are combined by passing them through a linear layer, which performs a matrix multiplication to produce an output that is 10 samples long. At this step, each sample represents the likelihood that the trace in question comes from a cell in the bursting mode, predicted at 1 second intervals.

The likelihood of each time step having come from the tonic state can then be inferred (Fig. 5B, grayscale boxes on the right). The state with the larger likelihood score is taken to be the final classification at each time step.

Five networks were produced, one for each fold of cross-validation. The median accuracy was 80.15% for the cross-validation phase and 82.02% for the test phase (Fig. 5C). The median F1-score was 0.7302 for the cross-validation phase and 0.8071 for the test phase (Fig. 5C). Next, we looked at the Receiver Operating Characteristic (ROC) curve for each network fold. This curve plots the false positive rate against the true positive rate of the classifier for various classification threshold values, and the area under this curve is an indicator of the quality of the classifier. The area under the ROC curve (AUC) for each of the five network folds is plotted in Fig 5E. The AUC values for all folds on both cross-validation and test data are much higher than 0.5, indicating that these networks have been trained well and make accurate predictions.

Of all the network folds, Fold 2 looked particularly promising, because it (1) consistently outperformed the other folds with respect to classification accuracy and F1-scores (Fig 5C, D, purple), and (2) had higher AUC values than the other network folds (Fig 5E,F). Thus, we decided to retain this network for future use, and have dubbed it CaMLsort (**Ca**lcium imaging and **M**achine **L**earning based tool to **sort** intracellular state).

### CaMLsort performs well on previously unseen calcium signal reconstructions

To be sure that networks weren’t performing well just because of peculiarities in the dataset used for training and testing, we presented an independent set of calcium signal reconstructions to the network that were not used in any phase of the training. These reconstructions (n=19) also had state switches in them. The ground truth class labels for each of the 19 cells (Fig. 6A, top) were compared with the raw predictions from the network (Fig. 6A, middle).

At first glance, it appears that the network isn’t making accurate calls, because each cell’s predicted state alternates between both states. However, these fluctuations are transient and the overall F1-scores for the raw predictions are >0.5 for 13 out of the 19 cells (median score 0.777). The distribution of F1-scores is shown in Fig. 6C. On closer observation, it is clear that the 6 cells for which F1-scores were low (< 0.5) had a switch in state during the recording. The poor prediction accuracy for these can be explained by the fact that the training dataset didn’t have any cases with state switches.

Given our prior knowledge that the state of a neuron doesn’t flicker rapidly between tonic and bursting, we decided to smoothen out the raw predictions using a majority voting strategy. For this, we took a rolling window of length *N* around each time step of the prediction and asked what the majority of the network calls were across all the windows that included the time step. If the average posterior scores for the majority class were higher than the average posterior scores for the minority class, we would confidently assign the majority class to that time step. If not, no confident call would be made for that time step. For example, to make a confident call at time t0 with a window length of 5, we first extract all the 5-step windows that include time t0 (there will be 5 such windows, except at the start and end, where there will be fewer). Next, we average the posterior scores within each of the windows and assign the ‘average state’ as the state with the higher posterior score. In this case, let’s say these were 2:3 in favour of bursting. If the average bursting posterior score for the 3 “bursting” windows was greater than the average tonic posterior score for the 2 “tonic” windows, the state for time t0 would be confidently assigned as “bursting”. If not, the prediction at time t0 would be left blank.

Using this majority voting strategy with a window length of 7, we were able to make confident calls for >90% of the duration for all 19 cells. As seen in Fig. 6B, the jitter in the raw CaMLsort predictions was considerably smoothened out by this process, and the F1-scores showed a marginal improvement as well, with a median F1-score of 0.8194 (Fig. 6C). As before, the cases where F1-scores were <0.5 corresponded to recordings with state switches. As an extra control, we obtained CaMLsort predictions from signals that were scrambled. These predictions had much lower F1- scores (Fig. 6C). In fact, when presented with scrambled traces, CaMLsort almost always (99.76% of the time) predicted the state to be tonic, with a very large, consistent differential between the posterior scores for the tonic (mean ± stdev: 0.7810 ± 0.0565) and bursting (mean ± stdev: 0.2190 ± 0.0565) classes. This implies that CaMLsort has, indeed, captured a latent trend present only in the true signal and not in a collection of random numbers of comparable magnitude.

### CaMLsort performs well on calcium signal reconstructions from simulated spike trains

As a further challenge to the trained networks, we decided to generate artificial spike trains that represent canonical cases of the tonic or bursting class, reconstruct the calcium signal for these spike trains and ask the network to classify these completely synthetic data (Fig. 7). We simulated Purkinje neuron simple spiking using either a Poisson model (tonic state) or by randomly sampling from the distribution of inter-event intervals from experiments (bursting state; also see Methods). The CF input spike train was considered to be independent and was generated using a Poisson model.The ground truth states for these simulated spike trains is shown in Fig. 7A (top trace). The raw CaMLsort predictions obtained for these data had a median F1-score 0.8571 (Fig. 7A, middle trace and 7C), although these too, had several reversals in the state predictions. On applying the majority voting strategy described above, however, these reversals were smoothened out (Fig. 7B), yielding a median F1-score of 0.9189 (Fig. 7C) and a confident prediction for at least 80% of the recording duration in every cell. As was observed in the previous case, scrambling the signal prior to prediction by the neural network were all of one class (tonic), yielding F1-scores which were either 1 (all right) or 0 (all wrong) (Fig. 7C).

### Neural networks trained only on simulated calcium signals correctly identify Purkinje neuron state from real data

Thus far, all the inputs and challenges to the network were made using calcium signal reconstructions. To test the applicability of these networks on data from experiments, we went back to our original simultaneous electrophysiology-imaging data and asked if the network could call the state of those cells correctly based on the dF/F traces alone.

We found that CaMLsort predicted the state of a cell from imaging data well (Fig. 8A), with a median F1-score of 0.7654 across all cells (Fig. 8C). This outcome is interesting, because the networks have only been exposed to simulated calcium signals generated using a first-order generative model in the training and test phases, and yet are able to reasonably assess the state from real data, which have several challenging features - variable sensor expression levels, shot noise and signal bleaching, to name a few.

Majority voting did not substantially improve the overall F1-score across all cells (median 0.75). However, by retaining only confident calls, the flip-flopping of network predictions was markedly reduced (Fig. 8B), as was observed before in the case of classifications using reconstructed calcium signals.

To benchmark the quality of these predictions, we obtained CaMLsort predictions from calcium signals reconstructed from the spike trains of these cells (since CaMLsort was trained only on simulated calcium signals). The prediction quality from reconstructions was comparable to that obtained from raw imaging data (median F1- score of 0.9011; Fig. 8C). Therefore, while CaMLsort is clearly able to classify cellular state from experimentally-obtained imaging data, its performance has room for improvement. Also consistent with our previous results were the predictions from scrambled dF/F traces, which were almost all (96.83%) tonic, thereby yielding F1- scores that were mostly either 0 or 1 (Fig. 8C).

To sum up, we showed that CaMLsort has learnt to sort the cellular state of a Purkinje neuron based only on its calcium signal and is capable of producing confident calls of high quality.

### CaMLsort generalizes well to calcium imaging data from other systems

Having developed a tool that can reasonably predict the bistable state of a Purkinje neuron given its calcium signal, we asked whether it could be broadly applicable to other neurons, too. Dopaminergic neurons of the ventral tegmental area (VTA) in mice also fire either tonically or phasically (Grace and Bunney, 1984a, b). Fleming et al., 2021 performed simultaneous electrophysiology and calcium imaging from these neurons, which expressed either GCaMP6f or GCaMP6m. This dataset is available publicly (https://github.com/jewellsean/spike_tools) and we tested CaMLsort on this dataset.

We first compiled the dataset and sorted cells as tonic or bursting, using a combination of visual inspection of spike rasters (Fig. 9A, B) and the coefficient of variation of inter-spike intervals (Fig. 9C), as we did for Purkinje neurons. Three independent reviewers used this method to classify states, and a consensus was taken across reviewers, while also taking into consideration the state labels provided by the authors in their metadata. Data were included only if at least 2 out of the 4 labels (3 reviewers plus original metadata) agreed. Using this method, 78/84 cells (92.86%) of the original dataset were included for further analysis. Of these, 16 cells were in the tonic state, while the remaining 62 cells were in the bursting state.

Before trying to use CaMLsort on this data, we asked whether the convolution-based forward model of reconstructing calcium signals from electrophysiology would work on this dataset. Since there is only a single kind of spike in these neurons, we used only one kernel, the amplitude, half-rise time constant and half-decay time constant of which were taken to be the mean values for GCaMP6f or GCaMP5G (Chen et al., 2013). No optimization of kernels was performed.

Visual inspection shows that reconstructions obtained from both GCaMP6f and GCaMP5G kernels were comparable to the signals imaged from VTA DA neurons, with the only difference between them being their amplitudes (Fig. 9D). To quantify the similarity between the ground truth and the two simulated calcium signals, we used the Pearson correlation coefficient. The correlation coefficients between the ground truth and reconstructions from both GCaMP6f and GCaMP5G kernels were nearly identical, with a value of 0.6400 ± 0.3189, on average, for both (Fig. 9E).

Reconstructions that were either scrambled or generated after shuffling the DA neuron’s spike times had no correlation with the ground truth signal, as expected (Fig. 9E).

Having confirmed that the Fleming et al. dataset had both tonic and bursting data, and that the mapping from electrophysiology to imaging was also possible with this data, we were ready to test CaMLsort on it. As before, we first interpolated the calcium signals to get a sampling frequency of 30Hz before inputting them to CaMLsort. Raw predictions from CaMLsort and their corresponding ground truth labels for a subset of 12 of the 78 cells are shown in Fig. 10A. The distribution of F1- scores obtained from all 78 cells, the median for which is 0.6841, is shown in Fig. 10B.

**Figure 10.**
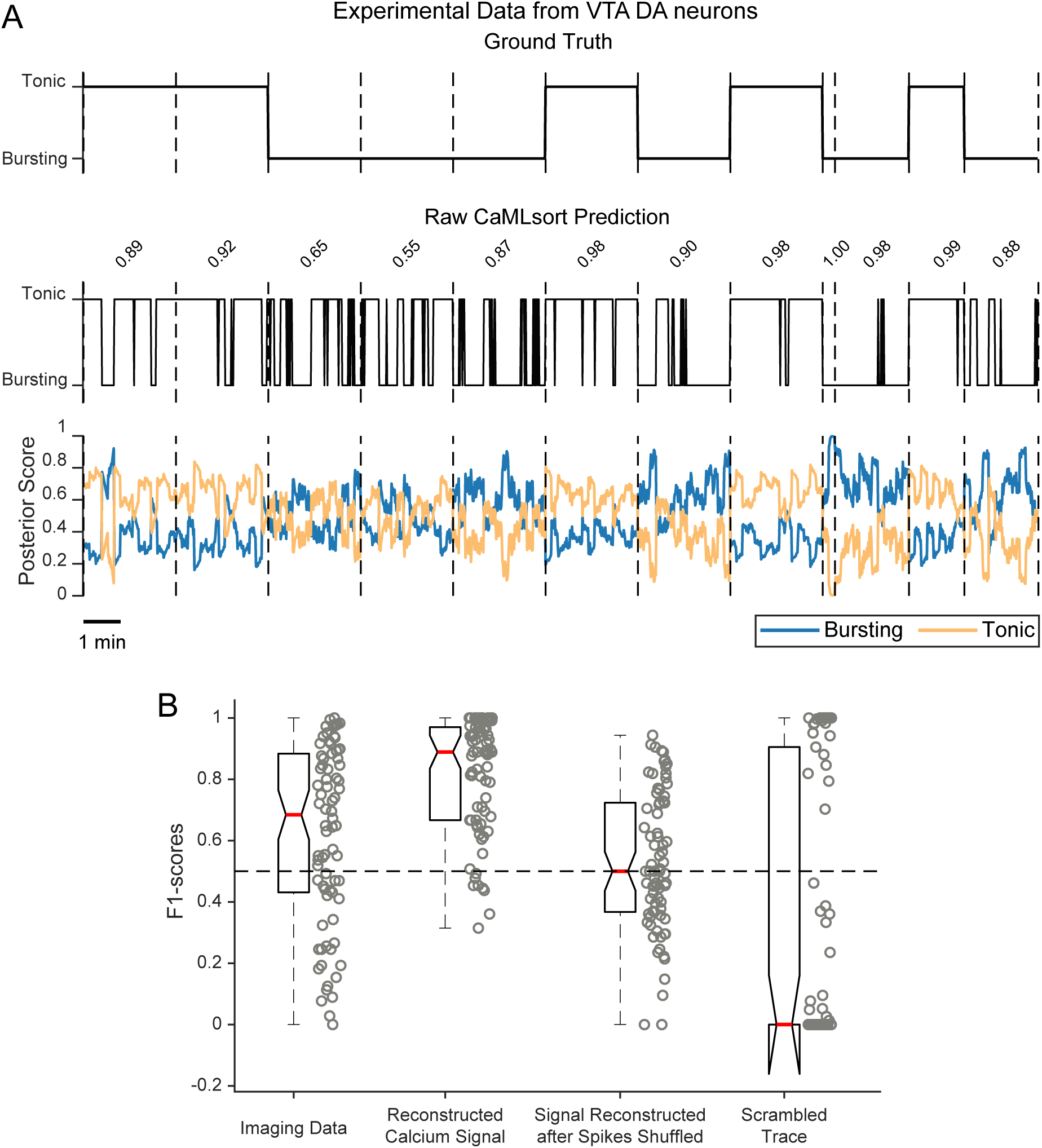
CaMLsort predictions of cellular state from calcium signals of VTA DA neurons (A) A subset of 12 (of 78) raw CaMLsort predictions from imaging data acquired from DA neurons of the VTA in mice. Each prediction was independently obtained but has been stitched together to produce a single time series with cells separated by vertical dashed lines. The ground truth state (top) is used as reference to assess the quality of CaMLsort’s predictions (middle) using the F1-score, which is printed above the prediction for each cell. The posterior scores for each class - bursting (blue) and tonic (light orange) - are shown in the lowest plot. **(B)** The entire distribution of F1-scores (n=78) between the ground truth state and the binary classification from CaMLsort obtained from various inputs - (from left to right) experimentally-acquired dF/F traces, calcium signals reconstructed from true spike trains using the GCaMP5G kernel, calcium signals reconstructed from shuffled spike trains using the GCaMP5G kernel, or a scrambled version of the raw dF/F data. Individual datapoints are shown alongside each boxplot. The horizontal dashed line corresponds to an F1-score of 0.5.

We also calculated F1-scores for predictions made on calcium signals reconstructed from true electrophysiology data using the GCaMP5G kernel (Fig. 10B). We decided to use the 5G kernel since the reconstructions that CaMLsort was trained on followed kinetics of GCaMP5G. The predictions from signal reconstructions were comparable to those from raw imaging data, with a median of 0.8889. Lastly, if the imaging signal was scrambled before prediction, CaMLsort predicted a default “tonic” state 89.03% of the time, yielding F1-scores that were mostly either 0 or 1 (Fig. 10B).

In all, our results from VTA DA neurons in mice were similar to those from zebrafish Purkinje neurons, which means that CaMLsort generalizes well and is likely to find broad use in classifying cellular state across systems. In order to facilitate this use, we have made CaMLsort publicly available as a GitHub package (https://github.com/vatsalat/CaMLsort23).

## Discussion

In this paper, we describe the development of CaMLsort, a machine learning-based tool to identify the bistable state of Purkinje neurons from calcium imaging data. We develop this network and train it only on a simulated calcium signal dataset, but find that it can be used on experimental data, too. Furthermore, we demonstrate that CaMLsort is able to classify bistable states in a completely different cell type (dopaminergic neurons) from a different brain region (ventral tegmental area) in a different model system (mice), and using a different version of the calcium sensor (GCaMP6) than it was originally trained on. While this demonstrates that CaMLsort could have wider application, more broader testing on other datasets will be essential to prove generalizability.

### Bistability in Purkinje neuron populations

The ability of neurons to exist in two stable states and exhibit distinct firing patterns/ excitabilities in those states has been documented in invertebrate (Dickinson and Nagy, 1983; Lechner et al., 1996; Marder et al., 1996; Russell and Hartline, 1982) and vertebrate nervous systems (Cowan and Wilson, 1994; Hounsgaard et al., 1984; Loewenstein et al., 2005; Misgeld et al., 1986; Schwindt and Crill, 1980).

We showed that larval zebrafish Purkinje neurons are bistable and that they switch between firing either tonically or in bursts (Sengupta and Thirumalai, 2015). Such behavior has also been reported in many other cell types such as thalamic (Jahnsen and Llinás, 1984) and cortical neurons (McCormick et al., 1985).

Though it is appreciated that bistability increases the information processing capacity of neurons, such as by allowing gating of synaptic inputs (Thirumalai and Jha, 2022), its implication for network computation or behavioural output is less well understood. This is true of Purkinje neurons as well - while it is established that these neurons are bistable and are important for motor behaviours (Engbers et al., 2013; Popa et al., 2018; Sengupta and Thirumalai, 2015), the role of bistability in Purkinje neuron computation or motor behaviour output has thus far not been clear.

We propose that having more than one stable state allows a cell to toggle in and out of participating in circuit computations. In the cerebellum, specifically, bistability may have significant consequences by modulating PN neuronal synchrony with other PNs. PNs make weak inhibitory synapses onto deep cerebellar nuclear (DCN) cells in mammals and eurydendroid cells in fish, with a high degree of convergence (Bae et al., 2009; De Zeeuw et al., 2011; Harmon et al., 2020; Ikenaga et al., 2006; Person and Raman, 2012). DCN cell spike times could be effectively controlled via inhibition from several PNs firing synchronously (Person and Raman, 2012). Since PNs fire tonically when synaptically-isolated and CF inputs can trigger bursts (Sengupta and Thirumalai, 2015), the bursting state is a network-embedded state while the tonic state is independent of network drive. By toggling between these two states PNs can increase or decrease the levels of population synchrony and thereby affect DCN firing differentially. Mapping state distribution across the PN population during various tasks will lead us to an understanding of how PNs synchronize or desynchronize during these tasks and how that in turn shapes cerebellar output and motor behaviors.

### Inferring long-term dynamics from calcium imaging data

Calcium imaging has been one of the core experimental techniques in neuroscience for more than three decades and has seen many major developments in sensor quality, microscopy and analytical algorithms and software (Chen et al., 2013; Göbel and Helmchen, 2007; Peron et al., 2015; Renninger and Orger, 2013; Tian et al., 2009; Yang and Yuste, 2017). Yet, the technique is largely used to identify regions of the brain that respond to presented stimuli or correlate with behavioral output under well-defined experimental conditions. This is resultant from the fact that the predominant mode of transient calcium elevation occurs due to the opening of voltage-gated calcium channels during action potential firing (Borst and Helmchen, 1998; Göbel and Helmchen, 2007). Thus, these calcium transients are quite brief, typically lasting only tens of milliseconds and report spiking activity of neuronal populations during sensory stimulation and/or motor output. However, it is clear that neurons also exhibit longer time scale dynamics such as bursting, which carries information for downstream neurons (Zeldenrust et al., 2018). As the currently available calcium sensors themselves are much slower compared to the calcium transients, it is obvious that the recorded dF/F signals must manifest signatures of such long-term dynamics. CaMLsort addresses this issue by looking at longer term signatures in the calcium imaging time series data to identify cellular state.

### Broader Applications of CaMLsort

The development of CaMLsort has opened up the possibility of a lot of future research in understanding PN bistability and its role in the function of the cerebellum as a whole. We can now experimentally address, for instance, what causes a switch in a PN’s state by doing a high-throughput screen for chemical agents that affect bistability. We know that while bistability is a cell-intrinsic property, state switches could be a result of modulation of network activity. Neuromodulators are key candidates for this, as they can alter cell-intrinsic properties as well as network activity, and could thereby affect the bistability of a neuron, as has been demonstrated in multiple experimental systems (Abbinanti et al., 2012; Conway et al., 1988; Hounsgaard et al., 1988; Lechner et al., 1996; Williams et al., 2002).

Another question we can now directly address is how the state of a cell changes over development. We hypothesize that the tonic state could be a developmentally early state, with the burst-like phenotype developing as they get integrated into the developing cerebellar circuit. By birth-dating PNs using a photoconvertible protein like Kaede and imaging their activity with GCaMP, we can use CaMLsort to ask whether younger cells tend to be more tonic or vice versa. While this question could potentially be addressed using electrophysiology, CaMLsort allows us to perform this experiment in a more high-throughput and minimally invasive fashion.

A number of cell types exhibit bistability including cortical neurons (Cowan and Wilson, 1994; Egorov et al., 2002), striatal neurons (Misgeld et al., 1986), thalamic neurons (Fuentealba et al., 2005) and spinal motoneurons (Hounsgaard et al., 1984). In all of these cases, cellular state dictates how synaptic inputs are integrated. Therefore, predicting cellular state is essential to understanding circuit computation and here CaMLsort will be immensely useful. It is natural to wonder whether CaMLsort will be equally effective in predicting cellular state in different cell types and in different model organisms. We have found that CaMLsort generalizes well to a completely different system than it was trained on - neurons from a different brain region (ventral tegmental area) in a different model organism (mouse) imaged using a later generation of GCaMP. We would caution against using CaMLsort right out of the box without validating its efficacy in the new system of interest using a small ground truth dataset.

Taken together, CaMLsort is a powerful tool that opens up new avenues of research into the functioning of any circuit with bistable neurons.

### Limitations of CaMLsort

Having said the above, CaMLsort is not free of limitations. For instance, the generative model we use to train the neural network is a linear, time-invariant model and results may improve with the use of more accurate models, such as ones that estimate the true increase in intracellular calcium concentration resulting from simple spikes and CF inputs. Furthermore, it could be that CaMLsort may not work independent of the imaging method used. In both the cases we tested CaMLsort, widefield epifluorescence microscopy was used. The signal and noise characteristics of other, more modern and popular, imaging methods like multiphoton and light sheet microscopy are different from traditional widefield imaging. However, we posit that this should also be easy to resolve, either by the acquisition of a small ground truth dataset, or by comparing the traces obtained from imaging to those in the reconstructions dataset, or by denoising the data prior to prediction using CaMLsort.

## Additional Information

### Data Availability

All data are available in the main text and figures. The CaMLsort package codes are available on Github: https://github.com/vatsalat/CaMLsort23

### Author Contributions

Conceptualization: AV and VT Investigation: AV, SU, MS, PKG and VT Analysis: AV, SU, MS, PKG and VT Writing: AV and VT

## Acknowledgement

The authors would also like to thank P.T. Jagadeesh for maintaining the fish facility and members of the Fishfolk lab at NCBS for their inputs at various stages of this project. In particular, we would like to thank Ms. Meha Jadhav and Ms. Shivangi Verma for their help with independently verifying parts of this manuscript. The authors would also like to thank Prof. Rohini Balakrishnan for initiating this collaboration.

## References

Abbinanti MD, Zhong G, Harris-Warrick RM. 2012. Postnatal emergence of serotonin-induced plateau potentials in commissural interneurons of themouse spinal cord. Journal of Neurophysiology 108:2191–2202. doi:10.1152/jn.00336.2012

Ahrens MB, Orger MB, Robson DN, Li JM, Keller PJ. 2013. Whole-brain functional imaging at cellular resolution using light-sheet microscopy. Nat Methods 10:413–420. doi:10.1038/nmeth.2434

Ali F, Kwan AC. 2020. Interpreting in vivo calcium signals from neuronal cell bodies, axons, and dendrites: a review. Neurophotonics 7:011402. doi:10.1117/1.NPh.7.1.011402

Armstrong DM, Edgley SA. 1984. Discharges of Purkinje cells in the paravermal part of the cerebellar anterior lobe during locomotion in the cat. J Physiol (Lond) 352:403–424.

Bae Y-K, Kani S, Shimizu T, Tanabe K, Nojima H, Kimura Y, Higashijima S, Hibi M. 2009. Anatomy of zebrafish cerebellum and screen for mutations affecting its development. Dev Biol 330:406–426. doi:10.1016/j.ydbio.2009.04.013

Borst JG, Helmchen F. 1998. Calcium influx during an action potential. Methods Enzymol 293:352–371. doi:10.1016/s0076-6879(98)93023-3

Chen T-W, Wardill TJ, Sun Y, Pulver SR, Renninger SL, Baohan A, Schreiter ER, Kerr RA, Orger MB, Jayaraman V, Looger LL, Svoboda K, Kim DS. 2013. Ultrasensitive fluorescent proteins for imaging neuronal activity. Nature 499:295–300. doi:10.1038/nature12354

Conway BA, Hultborn H, Kiehn O, Mintz I. 1988. Plateau potentials in alpha- motoneurones induced by intravenous injection of L-dopa and clonidine in the spinal cat. J Physiol (Lond) 405:369–384. doi:10.1113/jphysiol.1988.sp017337

Cowan RL, Wilson CJ. 1994. Spontaneous firing patterns and axonal projections of single corticostriatal neurons in the rat medial agranular cortex. Journal of Neurophysiology 71:17–32. doi:10.1152/jn.1994.71.1.17

De Zeeuw CI, Hoebeek FE, Bosman LWJ, Schonewille M, Witter L, Koekkoek SK. 2011. Spatiotemporal firing patterns in the cerebellum. Nature ReviewsNeuroscience 12:327–344. doi:10.1038/nrn3011

Dickinson PS, Nagy F. 1983. Control of a central pattern generator by an identified modulatory interneurone in crustacea. II. Induction and modification of plateau properties in pyloric neurones. Journal of Experimental Biology 105:59–82. doi:10.1242/jeb.105.1.59

Egorov AV, Hamam BN, Fransén E, Hasselmo ME, Alonso AA. 2002. Graded persistent activity in entorhinal cortex neurons. Nature 420:173–178. doi:10.1038/nature01171

Engbers JDT, Fernandez FR, Turner RW. 2013. Bistability in Purkinje neurons: Ups and downs in cerebellar research. *Neural Networks*, Computation in the Cerebellum 47:18–31. doi:10.1016/j.neunet.2012.09.006

Fleming W, Jewell S, Engelhard B, Witten DM, Witten IB (2021) Inferring spikes from calcium imaging in dopamine neurons. PLOS ONE 16:e0252345.

Fuentealba P, Timofeev I, Bazhenov M, Sejnowski TJ, Steriade M. 2005. Membrane Bistability in Thalamic Reticular Neurons During Spindle Oscillations. Journal of Neurophysiology 93:294–304. doi:10.1152/jn.00552.2004

Göbel W, Helmchen F. 2007. In Vivo Calcium Imaging of Neural Network Function. Physiology 22:358–365. doi:10.1152/physiol.00032.2007

Grewe BF, Helmchen F. 2009. Optical probing of neuronal ensemble activity. *Current Opinion in Neurobiology*, Neuronal and glial cell biology ● New technologies 19:520–529. doi:10.1016/j.conb.2009.09.003

Grienberger C, Konnerth A. 2012. Imaging Calcium in Neurons. Neuron 73:862–885. doi:10.1016/j.neuron.2012.02.011

Harmon TC, McLean DL, Raman IM. 2020. Integration of Swimming-Related Synaptic Excitation and Inhibition by olig2+ Eurydendroid Neurons in Larval Zebrafish Cerebellum. J Neurosci 40:3063–3074. doi:10.1523/JNEUROSCI.2322-19.2020

He K, Zhang X, Ren S, Sun J. 2016. Deep Residual Learning for Image Recognition2016 IEEE Conference on Computer Vision and Pattern Recognition (CVPR). Presented at the 2016 IEEE Conference on ComputerVision and Pattern Recognition (CVPR). pp. 770–778. doi:10.1109/CVPR.2016.90

Hochreiter S, Schmidhuber J. 1997. Long Short-Term Memory. Neural Computation 9:1735–1780. doi:10.1162/neco.1997.9.8.1735

Hounsgaard J, Hultborn H, Jespersen B, Kiehn O. 1988. Bistability of alpha- motoneurones in the decerebrate cat and in the acute spinal cat after intravenous 5-hydroxytryptophan. The Journal of Physiology 405:345–367. doi:10.1113/jphysiol.1988.sp017336

Hounsgaard J, Hultborn H, Jespersen B, Kiehn O. 1984. Intrinsic membrane properties causing a bistable behaviour of alpha-motoneurones. Exp BrainRes 55:391–394. doi:10.1007/BF00237290

Ikenaga T, Yoshida M, Uematsu K. 2006. Cerebellar efferent neurons in teleost fish. Cerebellum 5:268–274. doi:10.1080/14734220600930588

Jahnsen H, Llinás R. 1984. Electrophysiological properties of guinea-pig thalamic neurones: an in vitro study. The Journal of Physiology 349:205–226. doi:10.1113/jphysiol.1984.sp015153

Kerr JND, Greenberg D, Helmchen F. 2005. Imaging input and output of neocortical networks in vivo. Proceedings of the National Academy of Sciences 102:14063– 14068. doi:10.1073/pnas.0506029102

Kingma DP, Ba J. 2017. Adam: A Method for Stochastic Optimization. doi:10.48550/arXiv.1412.6980

Kitamura K, Häusser M. 2011. Dendritic calcium signaling triggered by spontaneous and sensory-evoked climbing fiber input to cerebellar Purkinje cells in vivo. J Neurosci 31:10847–10858. doi:10.1523/JNEUROSCI.2525-10.2011

Knogler LD, Kist AM, Portugues R. 2019. Motor context dominates output from purkinje cell functional regions during reflexive visuomotor behaviours. eLife 8:e42138. doi:10.7554/eLife.42138

Kwan AC, Dan Y. 2012. Dissection of Cortical Microcircuits by Single-Neuron Stimulation In Vivo. Current Biology 22:1459–1467. doi:10.1016/j.cub.2012.06.007

Lechner HA, Baxter DA, Clark JW, Byrne JH. 1996. Bistability and its regulation by serotonin in the endogenously bursting neuron R15 in Aplysia. J Neurophysiol 75:957–962.

Lev-Ram V, Miyakawa H, Lasser-Ross N, Ross WN. 1992. Calcium transients in cerebellar Purkinje neurons evoked by intracellular stimulation. Journal of Neurophysiology 68:1167–1177. doi:10.1152/jn.1992.68.4.1167

Lin B-J, Chen T-W, Schild D. 2007. Cell type-specific relationships between spiking and [Ca2+]i in neurons of the Xenopus tadpole olfactory bulb. The Journal of Physiology 582:163–175. doi:10.1113/jphysiol.2006.125963

Loewenstein Y, Mahon S, Chadderton P, Kitamura K, Sompolinsky H, Yarom Y, Häusser M. 2005. Bistability of cerebellar Purkinje cells modulated by sensory stimulation. Nat Neurosci 8:202–211. doi:10.1038/nn1393

Marder E, Abbott LF, Turrigiano GG, Liu Z, Golowasch J. 1996. Memory from the dynamics of intrinsic membranecurrents. Proceedings of the National Academy of Sciences 93:13481–13486. doi:10.1073/pnas.93.24.13481

Markram H, Toledo-Rodriguez M, Wang Y, Gupta A, Silberberg G, Wu C. 2004. Interneurons of the neocortical inhibitory system. Nat Rev Neurosci 5:793–807. doi:10.1038/nrn1519

Matsui H, Namikawa K, Babaryka A, Köster RW. 2014. Functional regionalization of the teleost cerebellum analyzed in vivo. Proc Natl Acad Sci USA 111:11846– 11851. doi:10.1073/pnas.1403105111

McCormick DA, Connors BW, Lighthall JW, Prince DA. 1985. Comparative electrophysiology of pyramidal and sparsely spiny stellate neurons of the neocortex. Journal of Neurophysiology 54:782–806. doi:10.1152/jn.1985.54.4.782

Misgeld U, Calabresi P, Dodt HU. 1986. Muscarinic modulation of calcium dependent plateau potentials in rat neostriatal neurons. Pflugers Arch407:482–487. doi:10.1007/BF00657504

Miyakawa H, Lev-Ram V, Lasser-Ross N, Ross WN. 1992. Calcium transients evoked by climbing fiber and parallel fiber synaptic inputs in guinea pig cerebellar Purkinje neurons. J Neurophysiol 68:1178–1189. doi:10.1152/jn.1992.68.4.1178

Moreaux L, Laurent G. 2007. Estimating firing rates from calcium signals in locust projection neurons in vivo. Frontiers in Neural Circuits 1.

Narayanan S, Thirumalai V. 2019. Contributions of the Cerebellum for Predictive and Instructional Control of Movement. Curr Opin Physiol 8:146–151. doi:10.1016/j.cophys.2019.01.011

Paszke A, Gross S, Massa F, Lerer A, Bradbury J, Chanan G, Killeen T, Lin Z, Gimelshein N, Antiga L, Desmaison A, Kopf A, Yang E, DeVito Z, Raison M, Tejani A, Chilamkurthy S, Steiner B, Fang L, Bai J, Chintala S. 2019. PyTorch: An Imperative Style, High-Performance Deep Learning LibraryAdvances in Neural Information Processing Systems. CurranAssociates, Inc.

Pedregosa F, Varoquaux G, Gramfort A, Michel V, Thirion B, Grisel O, Blondel M, Prettenhofer P, Weiss R, Dubourg V, Vanderplas J, Passos A, Cournapeau D, Brucher M, Perrot M, Duchesnay É. 2011. Scikit-learn: Machine Learning in Python. Journal of Machine Learning Research 12:2825–2830.

Peron S, Chen T-W, Svoboda K. 2015. Comprehensive imaging of cortical networks. *Current Opinion in Neurobiology*, Large-Scale Recording Technology (32) **32**:115–123. doi:10.1016/j.conb.2015.03.016

Person AL, Raman IM. 2012. Purkinje neuron synchrony elicits time-locked spikingin the cerebellar nuclei. Nature 481:502–505. doi:10.1038/nature10732

Popa LS, Streng ML, Ebner TJ. 2018. Purkinje Cell Representations of Behavior: Diary of a Busy Neuron. Neuroscientist 1073858418785628. doi:10.1177/1073858418785628

Ramirez JE, Stell BM. 2016. Calcium Imaging Reveals Coordinated Simple Spike Pauses in Populations of Cerebellar Purkinje Cells. Cell Reports 17:3125–3132. doi:10.1016/j.celrep.2016.11.075

Renninger SL, Orger MB. 2013. Two-photon imaging of neural population activity in zebrafish. *Methods*, Zebrafish Methods 62:255–267. doi:10.1016/j.ymeth.2013.05.016

Rupprecht P, Carta S, Hoffmann A, Echizen M, Blot A, Kwan AC, Dan Y, Hofer SB, Kitamura K, Helmchen F, Friedrich RW. 2021. A database and deep learning toolbox for noise-optimized, generalized spike inference from calcium imaging. Nat Neurosci 24:1324–1337. doi:10.1038/s41593-021-00895-5

Russell DF, Hartline DK. 1982. Slow active potentials and bursting motor patterns in pyloric network of the lobster, Panulirus interruptus. Journal of Neurophysiology 48:914–937. doi:10.1152/jn.1982.48.4.914

Schindelin J, Arganda-Carreras I, Frise E, Kaynig V, Longair M, Pietzsch T, Preibisch S, Rueden C, Saalfeld S, Schmid B, Tinevez J-Y, White DJ, Hartenstein V, Eliceiri K, Tomancak P, Cardona A. 2012. Fiji: an open-source platform for biological-image analysis. Nat Methods 9:676–682. doi:10.1038/nmeth.2019

Schwindt P, Crill W. 1980. Role of a persistent inward current in motoneuron bursting during spinal seizures. Journal of Neurophysiology 43:1296–1318. doi:10.1152/jn.1980.43.5.1296

Sengupta M. 2015. Purkinje neurons in action: From single cells to ensembles. Mumbai: Tata Institute of Fundamental Research.

Sengupta M, Thirumalai V. 2015. AMPA receptor mediated synaptic excitation drives state-dependent bursting in Purkinje neurons of zebrafish larvae. Elife 4. doi:10.7554/eLife.09158

Smetters D, Majewska A, Yuste R (1999) Detecting action potentials in neuronal populations with calcium imaging. Methods San Diego Calif 18:215–221.

Thirumalai V, Jha U. 2022. RECRUITMENT OF MOTONEURONS Vertebrate Motoneurons, In Press. Springer-Verlag.

Tian L, Hires SA, Mao T, Huber D, Chiappe ME, Chalasani SH, Petreanu L, Akerboom J, McKinney SA, Schreiter ER, Bargmann CI, Jayaraman V, Svoboda K, Looger LL. 2009. Imaging neural activity in worms, flies and micewith improved GCaMP calcium indicators. Nat Methods 6:875–881. doi:10.1038/nmeth.1398

Urasaki A, Morvan G, Kawakami K. 2006. Functional dissection of the Tol2 transposable element identified the minimal cis-sequence and a highly repetitive sequence in the subterminal region essential for transposition.Genetics 174:639– 649. doi:10.1534/genetics.106.060244

Vogelstein JT, Packer AM, Machado TA, Sippy T, Babadi B, Yuste R, Paninski L. 2010. Fast Nonnegative Deconvolution for Spike Train Inference From Population Calcium Imaging. Journal of Neurophysiology 104:3691–3704. doi:10.1152/jn.01073.2009

Williams SR, Christensen SR, Stuart GJ, Häusser M. 2002. Membrane potential bistability is controlled by the hyperpolarization-activated current I(H) in rat cerebellar Purkinje neurons in vitro. J Physiol (Lond) 539:469–483.

Yaksi E, Friedrich RW. 2006. Reconstruction of firing rate changes across neuronal populations by temporally deconvolved Ca2+ imaging. Nat Methods 3:377– 383. doi:10.1038/nmeth874

Yang W, Yuste R. 2017. In vivo imaging of neural activity. Nat Methods 14:349–359. doi:10.1038/nmeth.4230

Zeldenrust F, Wadman WJ, Englitz B. 2018. Neural Coding With Bursts—Current State and Future Perspectives. Frontiers in Computational Neuroscience 12.

